# Perinatal Exposure to Organophosphate Flame Retardants Induces Sex- and Hormone-Dependent Alterations in Anxiety, Memory, Neurotransmitter Content, and Hippocampal Gene Expression

**DOI:** 10.64898/2026.01.21.700648

**Authors:** Kimberly Wiersielis, Kevin Moran, RidhaFatima Mukadam, Jarrett Early, Victoria Appel, Catherine Rojas, Ali Yasrebi, Thomas Degroat, Nadja Knox, Troy A. Roepke

## Abstract

Developmental exposure to organophosphate flame retardants (OPFRs) is a public health concern due to their endocrine-disrupting potential. We examined perinatal exposure to tris(1,3-dichloro-2-propyl) phosphate, triphenyl phosphate, and tricresyl phosphate in mice. Adult male and female offspring were assessed for memory and anxiety-like behavior. Dopamine and norepinephrine were quantified in the hippocampus and prefrontal cortex (PFC), and bulk RNA sequencing was conducted for the hippocampus. OPFR-treated females in high ovarian hormone states spent less time in the open field test (OFT) center, the Y-maze unknown arm, and with the displaced object in spatial object recognition (SOR) indicating increased anxiety-like behavior and impaired spatial memory. These females also illustrated improved memory on the short-term Barnes maze, and a trending improvement in the novel object recognition test. Females in low ovarian hormone states, demonstrated a trend in center OFT exploration. OPFR-treated males displayed disruption in memory in the SOR and the short- and long-term Barnes maze. Perinatal OPFR reduced hippocampal dopamine in males and altered prefrontal dopamine in females in a hormone-dependent manner. OPFR-treated females in high ovarian hormones states demonstrated a trending decrease in PFC norepinephrine. Perinatal OPFR treatment caused differential gene expression in 121 individual genes and alteration to functional modules related to RNA processing, cellular metabolism, and extracellular organization. Hormone status also affected gene OPFR-induced altered expression, with similarity between males and high ovarian hormone state females. Our findings suggest that perinatal OPFR exposure causes widespread, sex specific, and hormone dependent disruptions in behavior, neurochemistry, and gene expression in adulthood.

**Highlights:**

- Anxiety-like behavior in OPFR-treated females varied with ovarian hormone status
- High ovarian hormone OPFR females showed task-dependent changes in memory
- Males displayed impaired spatial memory following perinatal OPFR treatment
- Perinatal OPFR modifies hippocampal and prefrontal dopamine and norepinephrine
- OPFR treatment altered individual gene and functional gene module expression

## 1. Introduction

Widespread contamination by endocrine disrupting chemicals (EDCs) is a critical environmental and public health issue. These compounds interfere with hormonal signaling pathways, leading to detrimental effects on developmental, reproductive, neurological, and immune systems in both humans and animal models [1–4]. Rapid industrial growth and consumer product manufacturing have contributed to the pervasive presence of EDCs in ecosystems and everyday items, including plastics, pesticides, and flame retardants [5–7]. As a result, human exposure is nearly constant, raising urgent concerns about long-term health consequences [8].

Flame retardants (FRs) are a major class of EDC exposure due to their widespread use and propensity to leach into the environment. These chemicals are added to a variety of materials to reduce flammability and delay ignition, with the goal of increasing escape time in the event of a fire [9, 10]. However, the chemical components of FRs are not completely bound to the materials they are intended to protect, allowing them to migrate into indoor environments over time. This results in their accumulation in household dust, which serves as a significant route of unintentional human exposure, particularly via ingestion [11]. Notably, tris(1,3-dichloro-2-propyl) phosphate (TDCPP), a commonly used FR, has been found in household dust at levels nearing 2000 ng/g [12]. Biomonitoring studies have confirmed systemic exposure: bis(1,3-dichloroisopropyl) phosphate (BDCIPP), a metabolite of TDCPP, has been consistently detected in the urine of children across the U.S. and globally, with median concentrations ranging from ∼2.8 ng/mL in the U.S. [13] to 7.8 μg/g in Australian children [14]. FR compounds have also been detected in blood (e.g., BDE-153, 0.13–3.1 pmol/g) [15], breast milk (e.g., tris-2-butoxyethyl phosphate, 1.44 ± 0.789 ng/mL) [16], and workplace environments such as office dust (TDCPP, mean 6.06 μg/g) [17], highlighting the pervasive nature of human exposure.

Polybrominated diphenyl ethers (PBDEs) were widely used as flame retardants in commercial products from the 1960s until the early 2000s, when mounting evidence linked them to a range of adverse health effects. In response, both regulatory actions and voluntary industry shifts led to the gradual phase-out of PBDEs [18]. This transition created a demand for alternative flame retardants, spurring the increased use of organophosphate flame retardants (OPFRs) to maintain fire safety standards [12]. OPFRs include a chemically diverse group of compounds, such as chlorinated alkyl phosphates and non-halogenated aryl phosphates. Notable formulations include triphenyl phosphate (TPP), a component of the commonly used mixture Firemaster® 550, TDCPP, and tricresyl phosphate (TCP). Evidence suggests that OPFRs may disrupt endocrine signaling by interacting with nuclear receptors involved in neuroendocrine regulation and metabolic homeostasis, including estrogen receptor alpha (ERα) and peroxisome proliferator-activated receptor gamma (PPARγ) [19–21]. Today, OPFRs have surpassed PBDEs as the most frequently detected flame retardants in consumer products [22, 23].

Exposure to FRs during critical windows of development poses a significant risk to the developing fetus. Experimental studies in rodents demonstrate that FRs can cross the placental barrier, directly impacting fetal neurodevelopment [24, 25]. These findings raise concern that maternal FR exposure may induce long-lasting alterations in brain development and function, leading to persistent neurobiological and behavioral changes in offspring. One particular domain of FR vulnerability is anxiety-like behavior [26]. Human studies link perinatal exposure to FRs with elevated anxiety symptoms in children [27–29]. Complementary findings in animal models further elucidate the effects of such exposures on anxiety-like behaviors. Perinatal exposure to Firemaster® 550 alters anxiety-like phenotypes in a sex-specific manner during adulthood [24, 30]. Baldwin et al. [24] investigated developmental exposure to Firemaster® 550 from gestational day (GD) 9 to 18 and produced sex-specific anxiety-like behaviors dependent on behavior test in adult Wistar rats. Exposed adult males showed increased anxiety in the elevated plus maze, evidenced by reduced exploration of open arms; however, females instead exhibited anxiety-like behavior in the light-dark box, where they delayed entry into the illuminated chamber. Interestingly, Patisaul and colleuges [30] noted a sex difference for perinatal exposure to Firemaster® 550 on anxiety-like behavior. Exposure to Firemaster® 550 from GD 8 through weaning resulted in sex-specific anxiety alterations in adult Wistar rats, as assessed by the zero maze (ZM). Females exhibited increased anxiety-like behavior, evidenced by fewer open arm entries at a 100-μg dose, whereas males exposed to the 100-μg dose showed increased open-arm exploration suggesting reduced anxiety. Our prior research demonstrated that maternal exposure to a mixture of organophosphate flame retardants (TPP, TDCPP, and TCP) induces sex-specific effects on anxiety-like behavior in adulthood. Male offspring exhibited an anxiolytic-like phenotype on the EPM following exposure, whereas female behavior remained unaffected [31].

Emerging evidence from both human and animal studies indicates that exposure to FRs during critical developmental windows can impair cognitive function and memory [26]. In humans, higher levels of urinary metabolites of OPFRs, such as BDCIPP and diphenyl phosphate (DPHP), have been associated with reduced full-scale IQ and poorer working memory performance in 7 year old children [32]. Poor cognitive findings from maternal FR exposure also occur in rodent models. Hawkey et al. [33] demonstrated that perinatal TPP exposure (16 or 32 mg/kg) in rats impaired recognition memory, as assessed by the novel object recognition test. Complementary findings in rodent models from our own work have demonstrated that perinatal exposure to OPFR mixtures (TPP, TDCPP, and TCP) results in deficits in spatial learning and memory during adulthood. Female mice exposed to OPFRs and maintained on a high-fat diet exhibited a reduced percentage of time spent exploring the unknown arm of the Y-maze compared to vehicle-treated controls, indicating impaired spatial memory; in contrast, OPFR-exposed males showed no significant behavioral differences [34].

The hippocampus is critical for memory acquisition, integration, and storage. Prior work has shown that perinatal OPFR exposure in rats causes a reduction in dentate gyrus volume in males, and alterations in markers related to neurogeneration and cellular maturation in males and females [35]. Others have demonstrated that TPP exposure during development impairs synaptogenesis alongside cognitive deficits [36]. The neurobehavioral consequences of maternal exposure to organophosphate flame retardants (OPFRs) remain poorly understood. In this study, we examined the long-term effects of developmental exposure to an OPFR mixture of tris(1,3-dichloro-2-propyl) phosphate (TDCPP), triphenyl phosphate (TPP), and tricresyl phosphate (TCP). Offspring were assessed in adulthood for anxiety-like behavior and cognitive performance using a battery of behavioral tests, including the open field test (OFT), Y-maze, spatial object recognition (SOR), novel object recognition (NOR), and the Barnes maze. We also quantified dopamine and norepinephrine levels in the hippocampus and the prefrontal cortex (prelimbic and infralimbic cortices), and conducted RNA sequencing of hippocampal tissue in male and female offspring with females stratified by ovarian hormone status (high: proestrus/estrus; low: metestrus/diestrus).

## 2. Methods

### 2.1. Animals and housing conditions

Adult male and female wild-type C57BL/6J mice (>60 days old) were bred in-house and housed in standard laboratory conditions. Animals were kept in conventional cages with unrestricted access to water and food (LabDiet PicoLab Verified 5v75 IF, <75 ppm phytoestrogens; Lab Diet, St. Louis, MO, USA). Environmental conditions were maintained at a temperature of 21–23 °C and humidity between 30–70%, with a 12-hour light/dark cycle (lights on at 0800 h). Mice were weaned at 3 weeks of age and pair housed by sex and litter. Weaned mice were moved to a reverse light cycle room 12-hour light, 12-hour dark cycle; lights off from 7:00 AM to 7:00 PM; controlled temperature (21–23 °C) and humidity (30–70%) regulation. Behavior testing on offspring began at 8 weeks. All procedures involving animals were conducted in accordance with institutional regulations aligned with National Institutes of Health guidelines and were approved by the Institutional Animal Care and Use Committee at Rutgers University.

### 2.2. Chemicals

The OPFR mixture was composed of tris(1,3-dichloro-2-propyl)phosphate (TDCPP, Sigma Aldrich, St. Louis, MO, CAS#: 13674-87-8, 95.6%), tricresyl phosphate (TCP, AccuStandard, New Haven, CT, CAS#: 1330-78-5, 99%), and triphenyl phosphate (TPP, Sigma Aldrich, St. Louis, MO, CAS#: 115-86-6, 99%), each administered at a dose of 1 mg/kg in an acetone:sesame oil vehicle (1:100). Similar dosing regimens have been used in previous perinatal OPFR exposure studies in rodents by our group and others [24, 30, 31, 37–41]. To prepare the OPFR stock solution, each compound was initially dissolved in 1 ml of acetone. Then, 100 μl of the acetone-based stock was diluted into 10 ml of non-estrogenic sesame oil and allowed to vent for 24–48 hours prior to use [37].

### 2.3. Maternal Exposures

Maternal exposure were completed as previously described [31, 40, 41]. Briefly, female baseline body weights were recorded throughout gestation, with GD 0 designated by the presence of a confirmed vaginal plug. Upon verifying pregnancy through sufficient weight gain, all mice were acclimated on GD 3 to oral administration using a mixture of dehydrated peanut butter and sesame oil (∼30 μl sesame oil). Male breeders were removed on GD 7, and pregnant females were then singly housed. Dams were randomly assigned to receive either the OPFR mixture or the vehicle control by oral administration from GD 7 through PND 14. This exposure window was selected to avoid disrupting implantation or early embryonic development and to encompass the hormonally sensitive period spanning birth through lactation. Dosing took place daily at 1000 h. Throughout the treatment phase, dams were housed individually; they remained singly housed during lactation. Offspring were weaned on PND 21 and subsequently housed by sex and litter, with 2 pups per cage.

### 2.4. Vaginal Cytology

Vaginal cytology was conducted following established methods to determine the stage of the estrous cycle[42, 43]. Lavage was performed on female mice following each behavioral test and on the day of tissue collection. The procedure involved gently flushing the vaginal canal with water, which was then transferred to a glass slide and evenly spread to form a monolayer. Slides were examined under a light microscope to assess the cellular composition and identify the specific stage of the estrous cycle based on characteristic cell types[44]. For analysis, females were grouped into two hormonal categories[45, 46]. Here, we grouped proestrus and estrus females, which correspond to periods of high ovarian hormone levels, and metestrus and diestrus females, which correspond to periods of low ovarian hormone levels.

### 2.5. Behavioral Analysis

All testing was recorded (ANY-maze, Stoelting, USA) by a camera suspended above the test arena. Mice went through a one month behavioral paradigm. Behavioral testing occurred between 10am and 2pm and was conducted during the dark phase of the light/dark cycle. All arenas were cleaned between trials and subjects. All behavior testing was conducted by an experimenter blind to experimental conditions.

#### 2.5.1. Y-maze

On the first day of behavioral testing, mice were evaluated using the Y-maze test, which consists of a white opaque plexiglass apparatus with three identical arms (8 cm wide, 15 cm high, 30 cm long) extending from a central triangular platform and designated as the start, habituation, and test arms. Opaque white removable doors were positioned 5 cm from the absolute center of the maze to block access to the arms during specific phases of the test. The maze was surrounded by outer walls, with fixed high-contrast visual cues (black-polka-dot) above the maze to provide consistent spatial orientation. The paradigm included two 5-minute phases: a habituation phase followed by a test phase[47]. During habituation, mice were placed at the distal end of the start arm and allowed to explore the start and habituation arms, while the test arm remained blocked. The assignment of habituation and test arms was counterbalanced across mice to control for side bias. Following habituation, mice were placed in a holding cage out of view of the testing arena for a 5-minute intertrial interval before being returned to the start arm and allowed to explore all three arms freely during the test phase. An arm entry was defined by all four paws entering an arm. Behavioral measures included: unknown arm time (sec), unknown arm entries, and the percentage of time (sec) and entries in the unknown arm.

#### 2.5.2 Open Field Test

On the second day of behavioral testing, mice began a four-day object location paradigm, which included an open field test (OFT) conducted on the first day as the habituation phase for spatial object recognition (SOR). The OFT was performed in a square, open-top arena constructed of white opaque plexiglass (40 cm × 40 cm × 40 cm) with a floor divided into a grid of 64 squares (8 × 8, each 5 cm per side). At the start of each trial, mice were placed in the same designated corner square (10 cm × 10 cm) and allowed to freely explore the arena for 5 minutes. All four paws needed to be in a zone to be classified as an entry. Automated behavioral analysis using ANY-maze software quantified several parameters, including time spent in the 10 cm center (sec) (defined as the central 2 × 2 grid squares), the percentage of time spent in the 10 cm center (sec), time spent in the 20 cm center (sec) (defined as the central 4 × 4 grid squares), the percentage of time spent in the 20 cm center (sec), total distance traveled (m), and mean speed (m/sec).

#### 2.5.3. Spatial Object Recognition (SOR)

From days 2 to 4 of behavioral testing, mice were habituated to the SOR arena in a white plexiglass square arena (40 cm × 40 cm × 40 cm, open top) containing a black and white vertical striped visual cue on one interior wall. Habituation sessions lasted 5 minutes per day over three consecutive days. No objects were present on habituation days. On the test day (Day 5), mice underwent a 5-minute training phase in which they were exposed to two identical plastic objects (50 ml conical tubes placed upside down) positioned at one end of the arena, each located 10 cm from the adjacent walls[48]. Objects were afixed to the arena with magnets. After training, mice were placed in a holding cage away from the SOR arena for a 5-minute delay period during which one of the objects was displaced to the opposite side of the arena. Following the delay, mice were returned to the arena for a 5-minute test phase, during which they were allowed to freely explore both objects. The initial training phase object placement and test phase object orientation of the displaced object were counterbalanced across subjects. Behavior was automatically scored using ANY-maze software, with object exploration defined as being within 2 cm of the object while actively sniffing. Mice that did not explore both objects during the test trial were excluded from analysis. Preference scores were calculated as the ratio of time spent with the displaced object to time spent with the familiar object. Behavioral measures included: the time spent investigating the displaced object (sec), number of interaction bouts with the displaced object, and the percentage of time (sec) and interaction bouts with the displaced object.

#### 2.5.4. Novel Object Recognition (NOR)

To confirm the novel object recognition (NOR) test was independent of hippocampal involvement, we employed a repeated habituation exposure training procedure previously shown to render the test hippocampus-independent [48–50]. Mice were exposed to the NOR paradigm following completion of SOR. Beginning the day after SOR testing, mice were re-habituated without objects to the same white plexiglass arena for 5 minutes per day over five consecutive days (Days 6 to 10). The test day (Day 11) followed a structure similar to SOR, with 5-minute training, delay, and test phases. During the training phase, mice were exposed to two identical metal objects (eyeglass holders; 5 cm × 5 cm × 5 cm). After a 5-minute delay period in a holding cage away from the arena, one of the metal objects was replaced with a novel glass object (upside down 125 mL flask; 11.5 cm × 6.5 cm × 6.5 cm) positioned in the same location. Mice were then returned to the arena for a 5-minute test phase and allowed to freely explore both objects. Initial location and object identity were counterbalanced across subjects. Behavior was automatically scored using ANY-maze software, with object exploration defined as being within 2 cm of the object while actively sniffing. Mice that did not explore both objects during the test trial were excluded from analysis. Preference scores were calculated as the ratio of time spent with the novel object to time spent with the familiar object. Dependent variables included time spent with the novel object (sec), number of interaction bouts with the novel object, and the percentage of time (sec) and interaction bouts with the novel object.

#### 2.5.5. Barnes Maze

Following the NOR test, the final behavioral assessment involved the Barnes maze. The platform mas made of white opaque plexiglass and measured 90 cm in diameter and contained 20 evenly spaced escape holes, each 5 cm in diameter and positioned 2 cm from the edge. The platform was elevated 60 cm above the ground by a cross-shaped base measuring 60 cm by 50 cm. During the training phase, a black opaque movable plexiglass escape tunnel (6 cm high, 20 cm long, 5 cm wide) with a 10 cm lip was secured beneath one of the escape holes by sliding it into ridges under the platform. The Barnes maze was placed in corner of the room, with fixed high-contrast visual cues (black-polka-dot) afixed to the wall. The Barnes maze paradigm was separated into a short-term memory test and a long-term memory test. For the short-term memory test paradigm, mice initially underwent ten training trials (2 per day) over five consecutive days (Days 12 to 16) on the elevated circular platform. At the beginning of the training trials, mice were placed in the center of the arena inside of a white opaque plexiglass start box (10 cm high × 10 cm wide; open bottom; moveable lid). The training trial was started by lifting the start box where the mice were allowed to freely explore the arena for 5 minutes to locate the escape tunnel. Between training trails mice were placed in their home cage. In an open environment, mice exhibit an innate tendency to seek out dark, enclosed spaces, a behavior typcially interpreted as an anxiety-like response driven by their instinct to avoid potential threats. The training trial was terminated early if the mice located the escape tunnel. Mice which could not located the escape tunnel were gently guided to it. During the training trails all mice were retained in the escape tunnel for 1 min. One hour after the last training trial mice were tested for short-term memory in the Barnes maze. For the test trial, mice were initally placed in the start box, upon its removal mice were allowed to freely explore the arena for a 5 min period. The escape tunnel was not present for the test trial. For the Barnes maze long-term memory, mice were tested one week later following the same testing paradigm. Automated behavioral analysis using ANY-maze software quantified several measures including: target quadrant time (sec), the number of target quadrant entries, time spent investigating the escape hole zone (sec), number of bouts into the escape hole zone, first entry escape hole distance (m), and escape hole zone latency (sec).

### 2.6. Dopamine and Norepinephrine ELISA

To quantify dopamine (DA) and norepinephrine (NE) levels, the prefrontal cortex (PFC) and dorsal hippocampus were dissected. Brains were sliced into 1 mm coronal sections using a brain matrix (Ted Pella, Redding, CA, USA), targeting the PFC (bregma +1.34 to +2.68 mm) (prelimbic and infralimbic corticies) and dorsal hippocampus (bregma -1.06 to -2.46 mm). Fresh sections were immediately microdissected under a dissecting microscope (Motic Microscopes), flash frozen, and stored at -80°C. During all stages of tissue handling, samples were kept on ice as frequently as possible to preserve sample integrity. For supernatant collection, tissue was homogenized in 40X buffer. Because tissue amounts varied slightly across samples due to manual dissection, the volume of buffer added was adjusted according to the weight of each tissue sample to ensure consistent homogenization. Samples were centrifuged at 1300 rpm for 10 minutes at 4°C. Supernatants were then transferred to fresh 1.7 mL microcentrifuge tubes and stored at -80°C for later ELISA analysis. DA and NE concentrations were quantified using ELISA kits specific for DA and NE (Rocky Mountain Diagnostics; catalogue #: BA E-5300R) (DA), and (Rocky Mountain Diagnostics; catalogue #: BA E-5200R) (NE). Absorbance was measured within 10 minutes using a microplate reader at 450, 620, and 650 nm wavelengths. The DA intra-assay coefficient of variation range was 10.8%–23.0% and NE intra-assay coefficient of variation range was 8.4%–15.6%.

### 2.7. Tissue Collection and RNA Sequencing

Tissue was collected and processed as previously described [42]. Mice were rapidly decapitated and brains were removed and sliced using a brain matrix into 1 mm blocks and the dorsal hippocampus was microdissect. Slices were then submerged in RNA-later solution (Invitrogen). Later the slices were microdissected isolating the dorsal hippocampus and tissue was again placed in RNAlater and stored at -80°C until RNA isolation. RNA was extracted using a RNAqueous Micro Isolation kit (Invitrogen; AM1931). RNA integrity was assessed with an Agilent 2100 Bioanalyzer. Any samples with an RNA integrity number (RIN) below 8 were excluded. The samples were then sent to the JP Sulzberger Columbia Genome Center (New York, NY) for sequencing. Library preparation was performed with the TruSeq Stranded mRNA Library Prep Kit (Illumina) and pooled libraries were sequenced on the Illumina NovaSeq 6000 with 100 bp paired-end reads at a 40 million read depth.

### 2.8. Bioinformatics Analysis

Raw RNA reads were processed using the Galaxy platform [51]. Briefly, the Trimmomatic tool was used to remove PCR duplicates and quality and adapter reads from Illumina sequencing, retaining reads with a minimum of 20 bases [52]. Processed reads were aligned to the mm10 reference genome using STAR and the NCBI RefSeq mm10 gene annotation [53].

Differential gene expression analysis was conducted using Bioconductor package limma [54]. For each treatment x hormone state condition, we conducted principal component analysis with filtered gene counts data (filtered out genes with less than 10 counts for each sample) and variance dendrogram analysis to visually inspect for outliers (Supplemental Fig. 1). One sample was removed from the oil x low ovarian hormone status group. Filtered read counts were then normalized to account for different library sizes among samples with a voom transform. Differentially expressed genes were identified between treatment groups with all subjects combined, between treatment group within hormone status group, and between hormone status group within treatment group. We adjusted raw p-value via empirical false discovery rate (eFDR) [55]. To estimate eFDR, we permuted sample IDs 5000 times (2500 times for within hormone status treatment comparisons due to smaller sample size) and obtained a null distribution of p-values. The significance threshold for differentially expressed genes (DEGs) was set as 15% of change in the absolute values of log 2-fold change (log2FC) at the eFDR of 5% (*p* < .05). Top significantly differentially expressed genes were selected by filtering these criteria and by lowest eFDR. We also performed gene ontology (GO) analysis to explore differences among identified DEGs within functional modules between groups using the clusterProfiler R package [56].

Weighted gene co-expression network analysis (WGCNA) was performed using the WGCNA R package [57]. WGCNA constructs Pearson correlation matrices of RNA expression and clusters highly co-expressed genes into several modules. WGCNA then examines the association between given experimental factors and module eigengenes (MEs), assigned to colors, which are the first component from the principal component analysis (PCA) for each module. WGCNA further calculates module membership for each gene. Module membership (MM) is measured as the Pearson correlation between the gene expression level and the module eigengene, with an absolute value of module membership close to 1 indicating that the gene is highly connected to other genes in the module. Gene counts were normalized with the DESeq2 package and genes that had low expression counts (<15) in more than 75% of samples were filtered out prior to network construction [58]. Each WGCNA model was constructed in signed hybrid mode. Power was selected by selecting a value with scale-free topology model fit above 0.8 with a minimal mean connectivity of 9. We tested whether each module eigengene was significantly different between groups using Pearson correlation and linear regression models, which is useful for identifying differential associations of MEs and has been used in other WGCNA analysis [59, 60]. Pearson correlation interaction are reported in WGCNA tables with (*R* = R value, and p values reported as * = *p* < .05, ** = *p* < .01). Results of linear models are presented as (*b* = beta coefficient ± standard error; n = number of observations in analysis; *p* = p value, two-tailed). We performed GO analysis on these modules to identify functions of independently clustered modules. Finally, we extracted hub genes with very high functional module membership. Finally, Welch’s t-tests were used to assess pairwise mean group differences for total gene expression, reported as [mean ± standard deviation; t(df) t value with estimated degrees of freedom; *p* = p value with α = .05, two-tailed, *d* = effect size using Cohen’s d].

### 2.9. Statistical Analysis

All behavior and ELISA data were analyzed with a 2 × 3 ANOVA with exposure (oil × OPFR) and hormonal status (male × female low ovarian hormones × female high ovarian hormones) as factors on Prism 10 (GraphPad Software, La Jolla, CA, USA). Post hoc analyses were performed using the Holm-Šidák correction for multiple comparisons, consistent with the analytical approach implemented previously [41, 42, 50, 61, 62]. Between 1 and 6 offspring were used per litter, and we used the average for each litter within each hormonal status as one datapoint. There were a total of six groups: oil:male; oil:female low; oil:female high; OPFR:male; OPFR:female low; and OPFR:female high. Values from individual mice that exceeded 2 standard deviations (SD) above or below the group mean were considered outliers and excluded. The results are reported as mean (± SEM) and results were considered statistically significant at *p* < .05. For analysis of RNA sequencing, all statistical analyses were completed using R v4.3.2 [63, 64]. Hormone status condition was subdivided into Males, Low for females in metestrus or diestrus, and High for females in estrus or proestrus.

## 3. Results

### 3.1. Y-Maze

On time spent in the unknown arm (sec) (Fig. 1A), we did not observe an effect of treatment or an interaction effect; however, we did observe an effect of hormonal status (*F*(2,62) = 3.492, *p* = .0366). OPFR-treated females in high ovarian hormone states exhibited a trending decrease in time spent in the unknown arm in contrast to control high females (*p* = .0932). Next, we evaluated the number of entries in the unknown arm (Fig. 1B). We did not find an effect of treatment, hormonal status, nor an interaction effect. The percentage of time spent in the unknown arm (sec) (Fig. 1C) indicated no effect of sex, treatment, or interaction, although we did observe a significant pairwise difference between OPFR- and control-treated high females. OPFR-treated high females displayed a reduction in the percentage of time in the unknown arm (sec) in contrast to controls (*p* = .0392). The percentage of entries in the unknown arm (Fig. 1D) resulted in no differences in treatment, hormonal status, or an interaction between treatment and hormonal status.

**Fig. 1.**
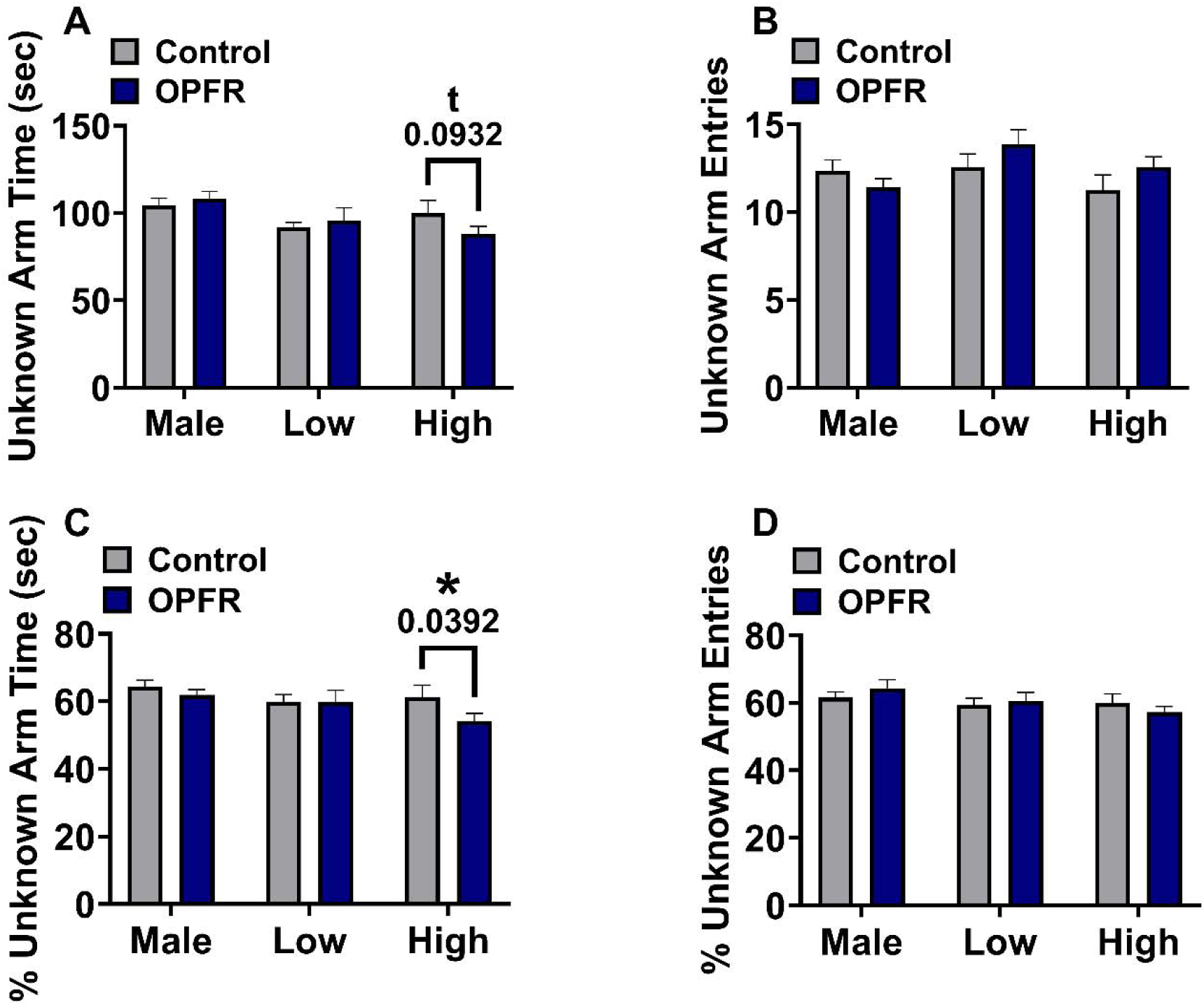
Y-Maze: OPFR-treated females in high (steroid) ovarian hormone states exhibited impaired hippocampal-dependent spatial navigation memory in the Y-maze. A) OPFR-treated high-steroid females spent a reduced amount of time in the unknown arm (sec) than their control counterparts. B) On unknown arm entries, no pairwise differences were observed. C) OPFR-treated high-steroid females demonstrated a reduction in the percentage of time in unknown arm (sec). D) No pairwise differences were observed on the percentage of entries in the unknown arm. Data are represented as mean ± SEM. * = *p* < .05; t = *p* < .10.

### 3.2. Open Field Test

On the amount of time spent in the 10cm center (sec) (Fig. 2A), we did not find an effect of treatment or hormonal status, but we did find an interaction effect between treatment and hormonal status (*F*(2,61) = 3.225, *p* = .0466). OPFR-treated females in high ovarian hormone states had a trending reduction in 10cm center time (sec) in contrast to control high females (*p* = .0651). OPFR-treated low females had a trending increase in 10cm center time (sec) in contrast to control low females (*p* = .0929). Next, the percentage of time spent in the 10cm center (sec) was evaluated (Fig. 2B). We did not observe an effect of treatment or hormonal status, but we did find an interaction effect between treatment and hormonal status (*F*(2,61) = 3.308, *p* = .0432). OPFR-treated females in high ovarian hormone states had a reduction in the percentage of time in the 10cm center (sec) in contrast to control high females (*p* = .0628). OPFR-treated low females had a trending increase in 10cm center time (sec) in contrast to control low females (*p* = .0877). For the amount of time spent in the 20cm center (sec) (Fig. 2C), we did not detect an effect of treatment or hormonal status, but we did find an interaction effect (*F*(2,62) = 3.592, *p* = .0334). OPFR-treated high-steroid females had a significant reduction in the amount of time spent in the 20cm center (sec) in contrast to control-treated high-steroid females (*p* = .0167). Considering percentage of time spent in the 20cm center (sec) (Fig. 2D), we found an interaction effect (*F*(2,62) = 3.746, *p* = .0291), with no differences in treatment or hormonal status. OPFR-treated high-steroid females had a reduction in the percentage of time in the 20cm center (sec) in contrast to control-treated high-steroid females (*p* = .0141). For both distance (m) (Fig. 2E) and mean speed (m/sec) (Fig. 2F), we did not find an effect of treatment, hormonal status, nor an interaction between treatment and hormonal status.

**Fig. 2.**
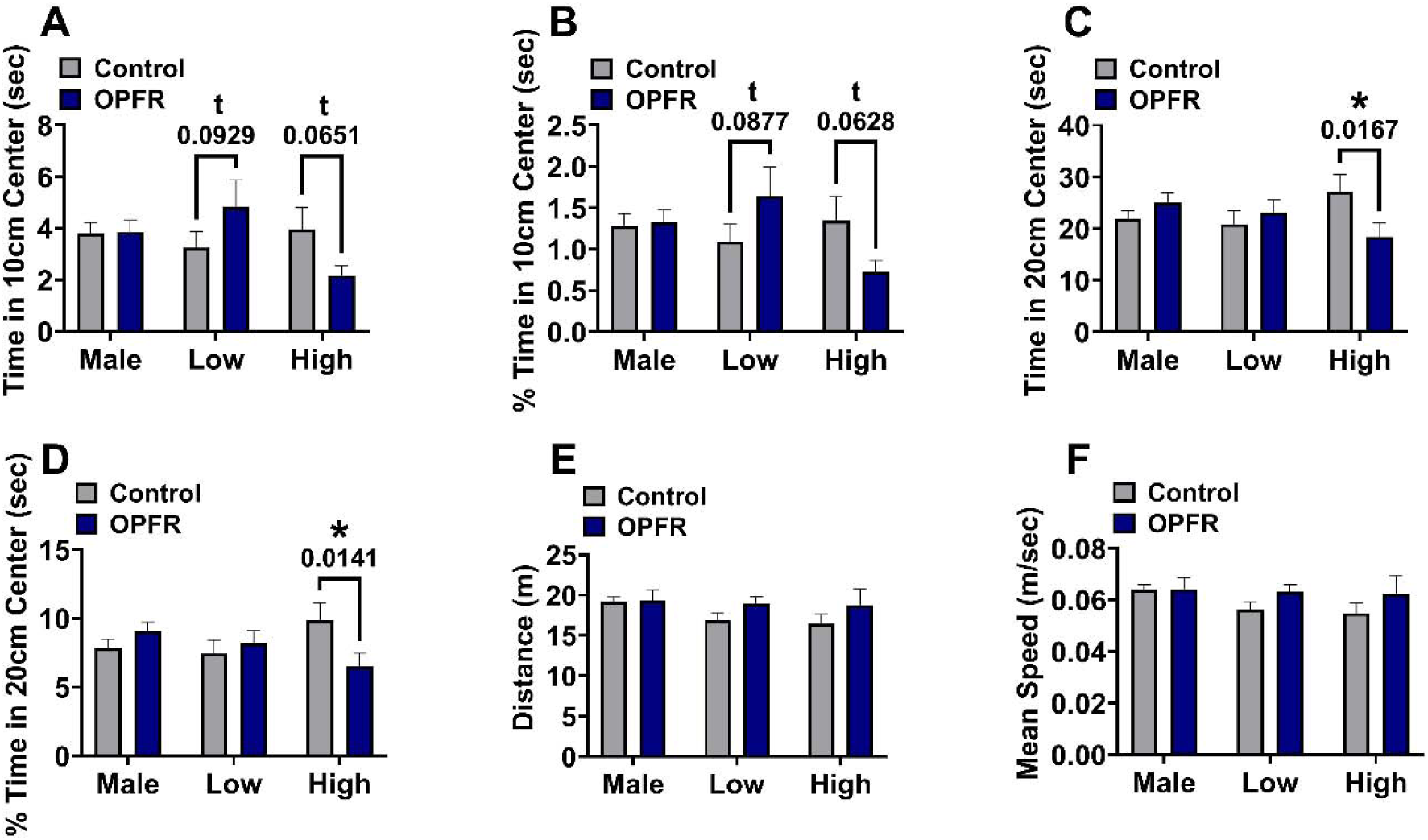
Open Field Test: OPFR-treated females in low ovarian hormone states demonstrated a trending anxiolytic-like response and OPFR-treated females in high ovarian hormone states displayed an anxiogenic-like response. A) OPFR-treated low-steroid females had a trending increase in time in the 10cm center (sec). OPFR-treated high-steroid females depicted a trending decrease. B) For the percentage of time in the 10cm center (sec), OPFR-treated low-steroid females demonstrated a trending increase, and OPFR-treated high-steroid females exhibited a trending reduction. C) OPFR-treated high-steroid females illustrated a significant decrease in time spent in the 20cm center zone (sec). Considering the percentage of time in the 20cm center (sec), OPFR-treated high-steroid females presented a significant reduction. E) Distance (m) revealed no differences between groups. F) We did not observe any differences in mean speed (m/sec). Data are represented as mean ± SEM. * = *p* < .05; t = *p* < .10.

### 3.3. Spatial Object Recognition (SOR)

The analysis of time spent with the displaced object (sec) (Fig. 3A) revealed no effect of treatment or hormonal status but did yield a trending interaction between treatment and hormonal status (*F*(2,57) = 2.816, *p* = .0682). OPFR-treated males had a trending reduction in time spent with the displaced object (sec) in contrast to control-treated males (*p* = .0903). For the number of displaced object bouts (Fig. 3B), we observed no effect of treatment or hormonal status; however, we did find a trending interaction between treatment and hormonal status (*F*(2,57) = 3.010, *p* = .0572). OPFR-treated males had a significant reduction in the number of bouts with the displaced object in contrast to control-treated males (*p* = .0498). The percentage of time spent with the displaced object (Fig. 3C) did not exhibit an effect of treatment, hormonal status, nor an interaction effect. We did detect a pairwise difference between OPFR-treated and control-treated high-steroid females. OPFR-treated females in high steroid states had a significant reduction in the percentage of time with the displaced object in contrast to control-treated high-steroid females (*p* = .0497). Lastly, for the percentage of displaced object bouts (Fig. 3D), we uncovered a significant effect of treatment (*F*(1,59) = 4.999, *p* = .0292), but no effect of hormonal status or interaction between treatment and hormonal status. OPFR-treated high-steroid females demonstrated a reduction in the percentage of displaced object bouts in contrast to high female controls (*p* = .0368). In addition, we observed a trending reduction in OPFR-treated males compared to control-treated males (*p* = .0656).

**Fig. 3.**
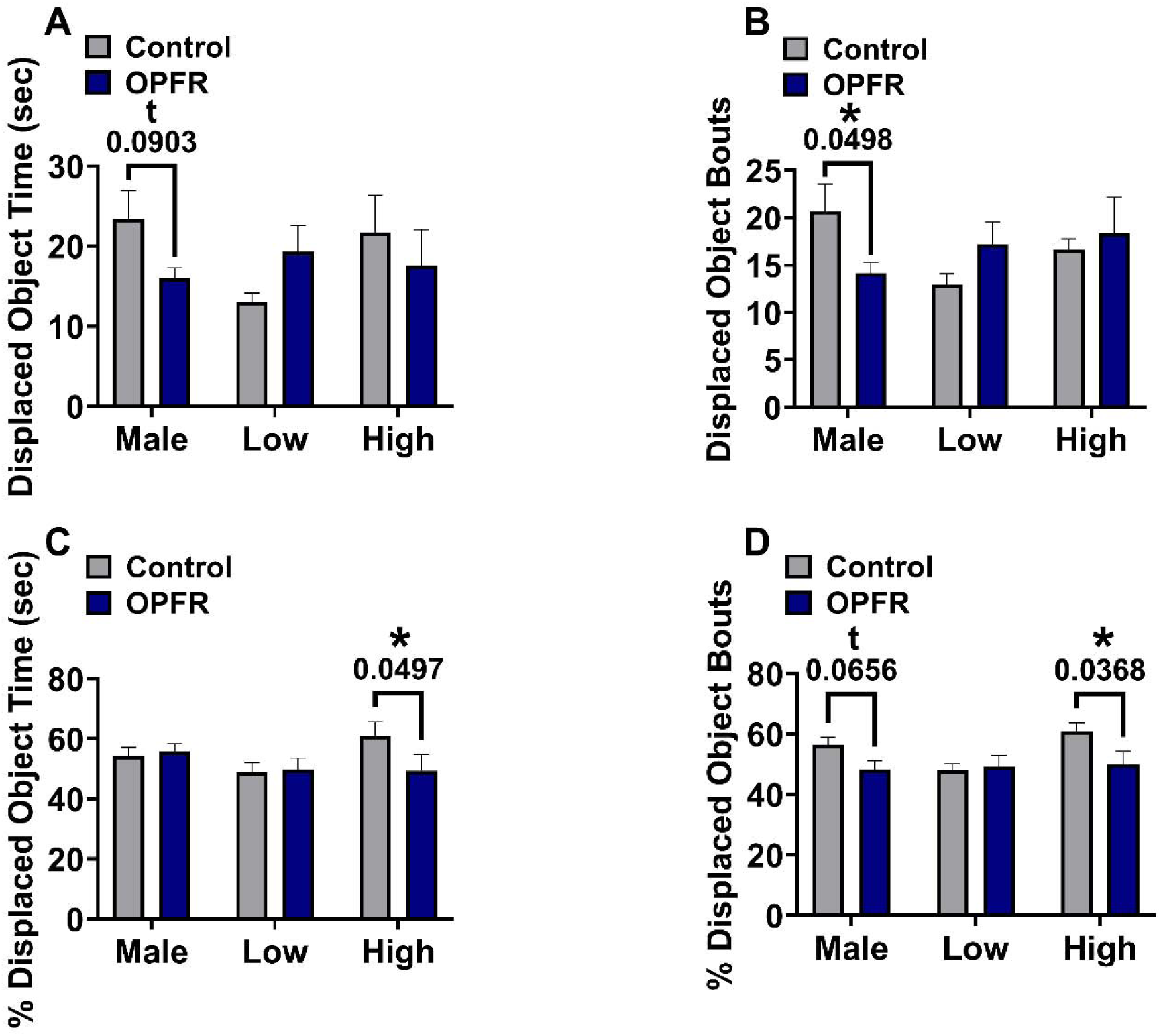
Spatial Object Recognition: OPFR-treated males and OPFR-treated high-steroid females showed a reduction in hippocampal-dependent spatial object orientation memory. A) OPFR-treated males displayed a trending reduction in time spent with the displaced object (sec). B) For the number of displaced object bouts OPFR-treated males illustrated a significant reduction. C) Considering the percentage of time spent with the displaced object (sec) OPFR-treated high-steroid females demonstrated a significant decrease. D) The percentage of displaced object bouts identified OPFR-treated males had a trending reduction and OPFR-treated females in high steroid states had a significant reduction in contrast to control-treated high-steroid females. Data are represented as mean ± SEM. * = *p* < .05; t = *p* < .10.

### 3.4. Novel Object Recognition (NOR)

On the analysis of time spent with the novel object (Fig. 4A), we did not observe an effect of treatment, hormonal status, or an interaction between treatment and hormonal status. The number of novel object bouts (Fig. 4B) demonstrated no effect of treatment, hormonal status, nor interaction effect. Pairwise comparisons revealed a trending difference between OPFR-treated females in high steroid states and control-treated high-steroid females, such that OPFR-treated females had an increase in novel object bouts in contrast to control-treated females (*p* = .0627). For the percentage of time spent with the novel object (Fig. 4C) and the percentage of novel object bouts (Fig. 4D), we did not find an effect of treatment, hormonal status, nor an interaction between treatment and hormonal status.

**Fig. 4.**
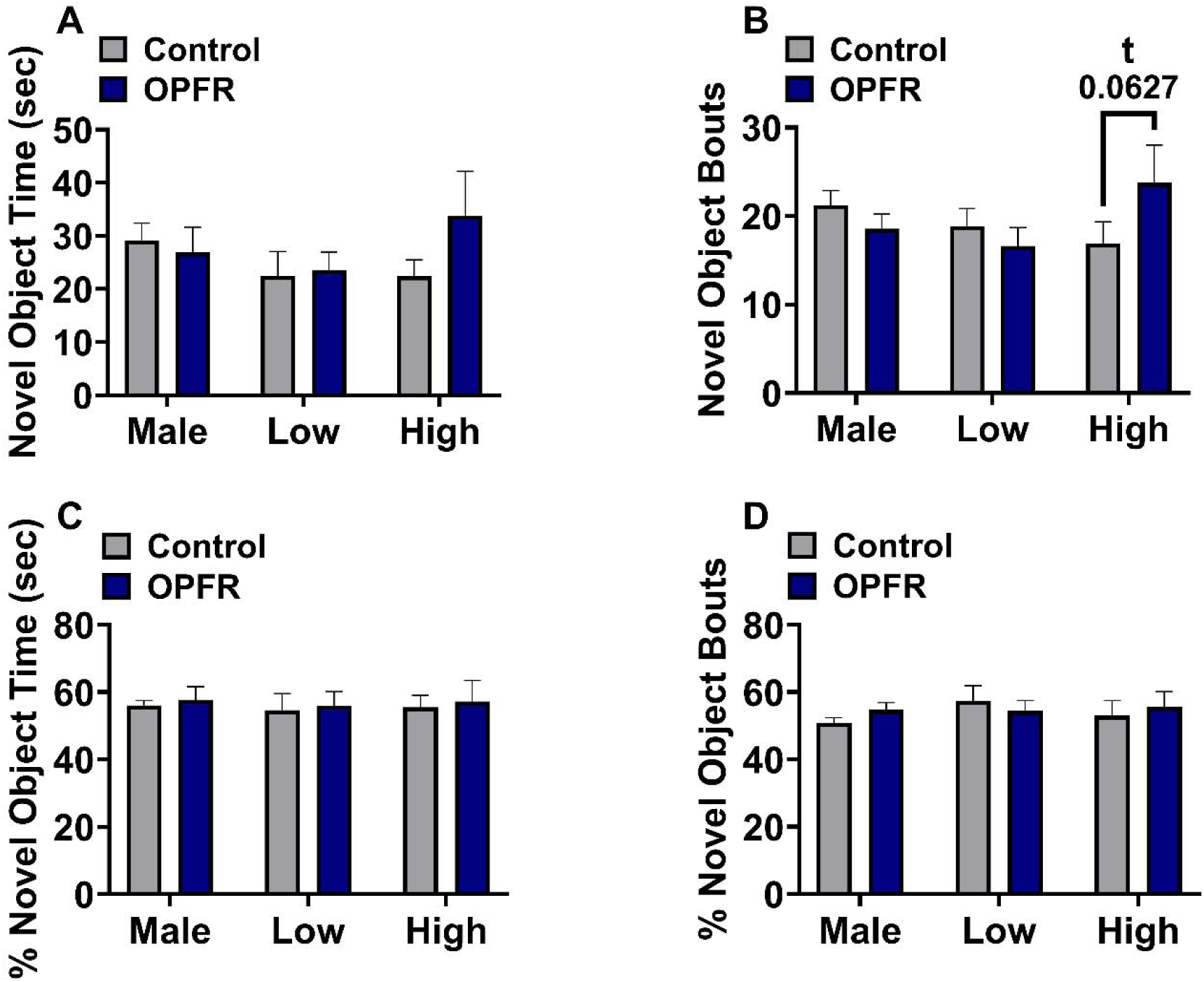
Novel Object Recognition: OPFR-treated females in high steroid states demonstrated a trending improvement in hippocampal-independent memory on the novel object recognition test. A) No differences were observed in novel object time (sec). B) OPFR-treated high-steroid females displayed a trending increase in the number of novel object bouts in contrast to control counterparts. C) For the percentage of time spent with the novel object (sec), no differences were detected. D) The percentage of novel object bouts did not demonstrate any pairwise differences. Data are represented as mean ± SEM. t = *p* < .10.

### 3.5. Short-term Memory Barnes Maze

The analysis of time spent in the target quadrant (sec) (Fig. 5A) revealed no effect of treatment or hormonal status but did yield a trending interaction between treatment and hormonal status (*F*(2,53) = 2.440, *p* = .0969). OPFR-treated males had a reduction in time spent in the target quadrant in contrast to control males (*p* = .0162). The number of target quadrant entries (Fig. 5B) uncovered no effect of treatment, hormonal status, nor an interaction effect. The amount of time spent with the escape hole (sec) (Fig. 5C), yielded no effect of hormonal status or an interaction effect; however, there was a trending effect of treatment (*F*(1,52) = 3.837, *p* = .0555). For the number of escape hole entries (Fig. 5D), we did not find an effect of treatment, but we did observe a trending effect of hormonal status (*F*(2,53) = 2.665, *p* = .0789) and a significant interaction between treatment and hormonal status (*F*(2,53) = 5.555, *p* = .0065). OPFR-treated females in high steroid states displayed an increased number of escape hole entries in contrast to control-treated high-steroid females (*p* = .0017). Next, for the first entry escape hole distance (m) (Fig. 5E), we did not find an effect of treatment, hormonal status, nor interaction between treatment and hormonal status. Lastly, for the escape hole latency (sec) (Fig. 5F), we did not find an effect of hormonal status; however, we did observe a trending effect of treatment (*F*(1,50) = 2.817, *p* = .0995) and a trending interaction effect (*F*(2,50) = 3.103, *p* = .0537). OPFR-treated females in high steroid states had a significant reduction in escape hole latency (sec) in contrast to control high females (*p* = .0101).

**Fig. 5.**
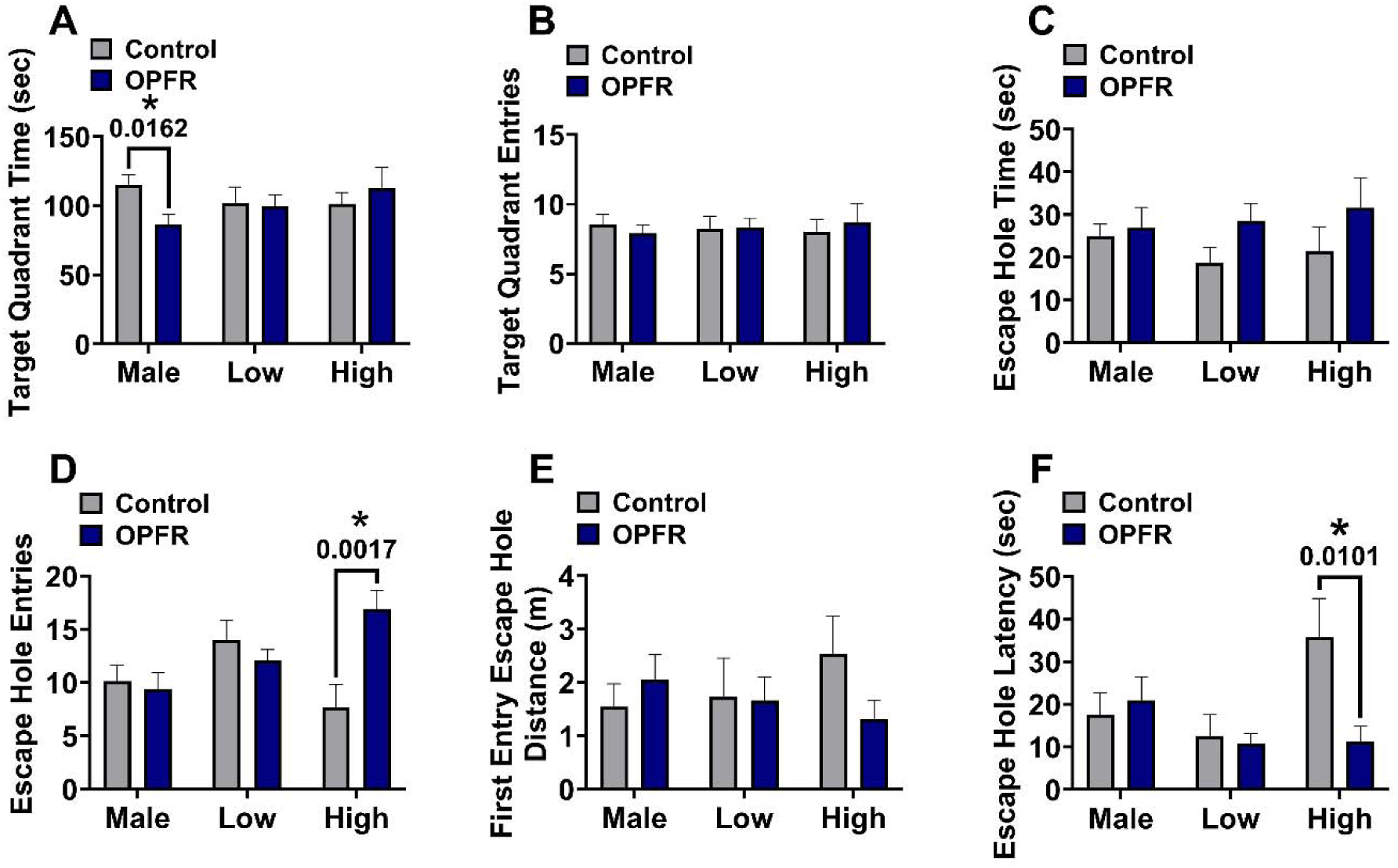
Short-term Memory Barnes Maze: OPFR-treated males exhibited an impairment in memory; although OPFR-treated high-steroid females displayed an improvement in memory. A) On the examination of target quadrant time (sec) OPFR-treated males had a reduction in contrast to control counterparts. B) No differences were observed on target quadrant entries. C) On the analysis of escape hole time (sec), no differences were detected. D) For escape hole entries, OPFR-treated high-steroid females illustrated a significant increase. E) For first entry escape hole distance (m), no pairwise differences were observed. F) OPFR-treated females in high steroid states showed a reduction in escape hole latency (sec) in contrast to control high females. Data are represented as mean ± SEM. * = *p* < .05.

### 3.6. Long-term Memory Barnes Maze

For the time spent in the target quadrant (sec) (Fig. 6A) we did not identify an effect of treatment, hormonal status, nor an interaction between treatment and hormonal status. Next, the number of target quadrant entries (Fig. 6B) revealed no effect of treatment or hormonal status; however, we did find a trending interaction effect (*F*(2,55) = 2.833, *p* = .0675). OPFR-treated males displayed a trending decrease in the number of target quadrant entries in contrast to control males (*p* = .0878). The amount of time spent with the escape hole (sec) (Fig. 6C), yielded no effect of treatment, hormonal status, or an interaction effect. For the number of escape hole entries (Fig. 6D) we did not observe an effect of treatment or hormonal status, but we did illustrate a significant interaction between treatment and hormonal status (*F*(2,55) = 3.835, *p* = .0276). OPFR-treated males displayed a decrease in the number of escape hole entries in contrast to control males (*p* = .0189). For both the first entry escape hole distance (m) (Fig. 6E) and escape hole latency (sec) (Fig. 6F), we did not reveal an effect of treatment, hormonal status, nor interaction between treatment and hormonal status.

**Fig 6.**
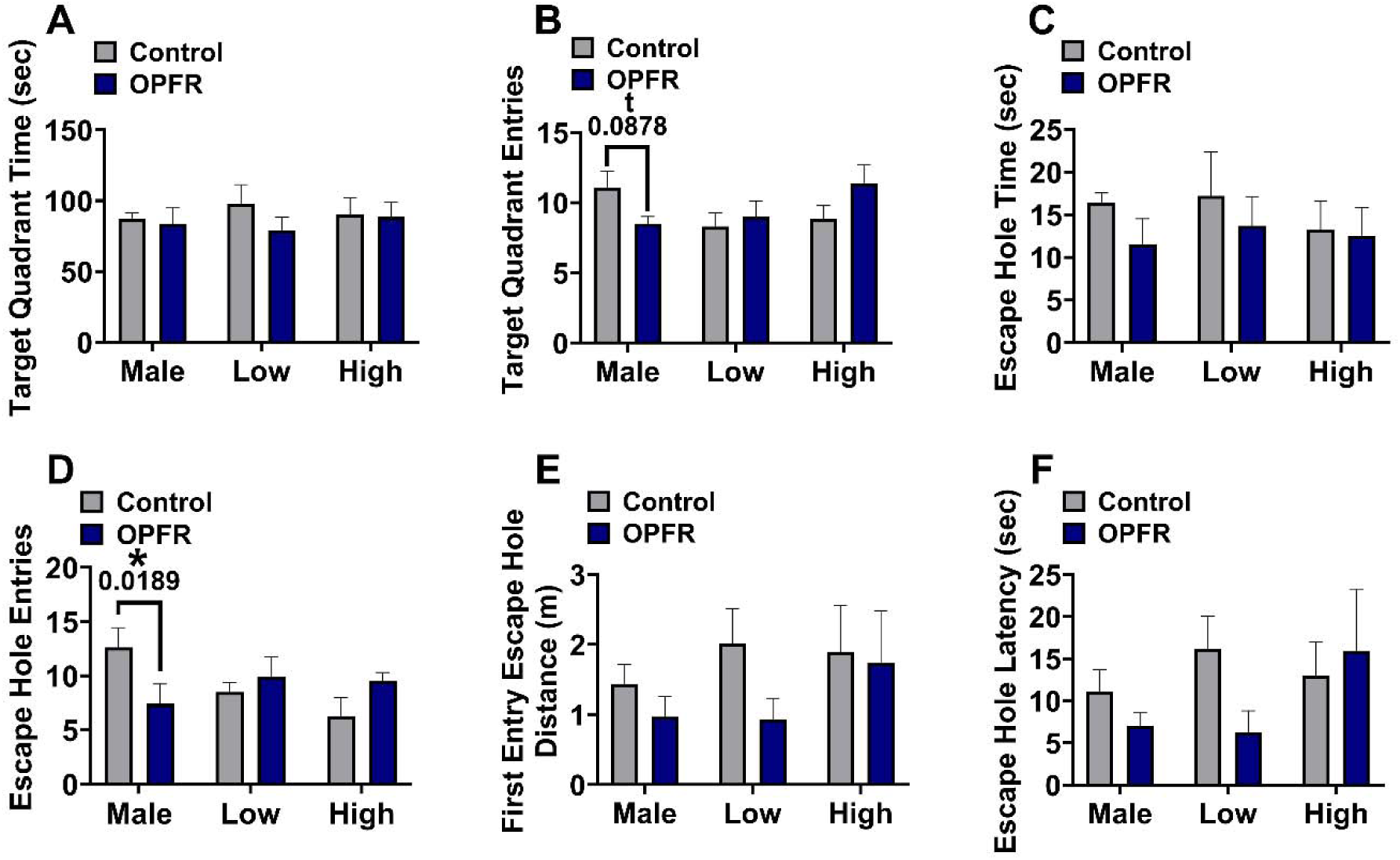
Long-term Barnes Maze: OPFR-treated males demonstrated an impairment in long-term memory. A) No differences were observed in the target quadrant time (sec). B) OPFR-treated males had a trending decrease in target quadrant entries. C) For escape hole time (sec) no differences were detected. D) On the assessment of escape hole entries, OPFR-treated males displayed a significant reduction in contrast to control males. E) For first entry escape hole distance (m), no pairwise differences were observed. F) On the analysis of escape hole latency (sec), no differences were demonstrated. Data are represented as mean ± SEM. * = *p* < .05; t = *p* < .10.

### 3.7. Neurotransmitters

#### 3.7.1 Dopamine

Considering the DA content in the dorsal hippocampus (Fig. 7A), we did not detect an effect of treatment or hormonal status, but we did reveal a trending interaction effect between treatment and hormonal status (*F*(2,51) = 2.804, *p* = .0700). OPFR-treated males had a decreased amount of DA content in the dorsal hippocampus in contrast to control-treated males (*p* = .0439). Next, examining DA content in the combined prelimbic and infralimbic cortices (Fig. 7B), we did not find an effect of treatment or hormonal status; however, we did demonstrate an interaction effect between treatment and hormonal status (*F*(2,51) = 3.914, *p* = .0262). OPFR-treated females in low steroid states had a trending reduction in DA content in contrast to control-treated low females (*p* < .0715). Moreover, OPFR-treated females in high steroid states displayed increased DA content in contrast to control-treated females in high steroid states (*p* = .0432).

**Fig. 7.**
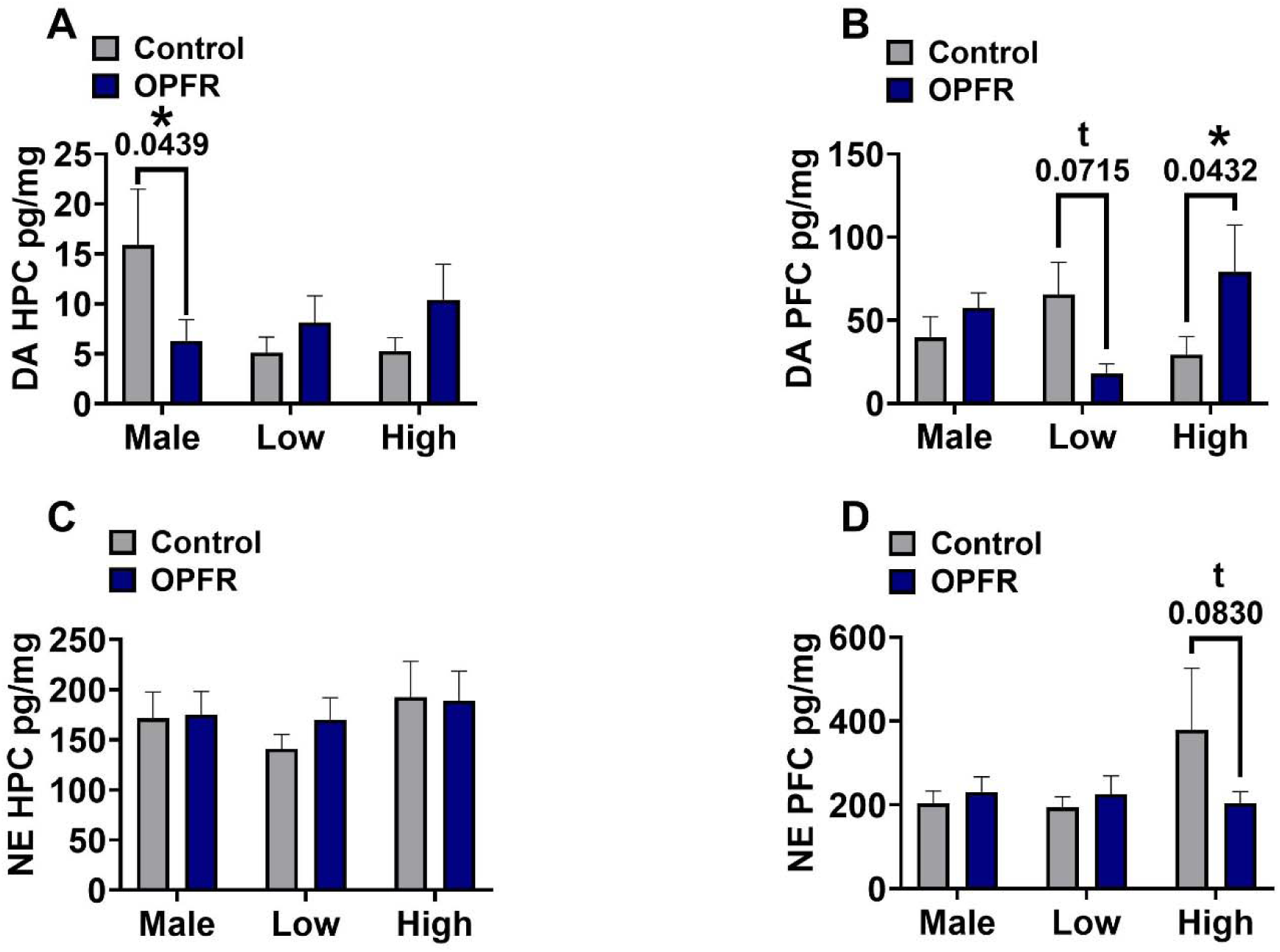
OPFR treatment altered dopamine (DA) and norepinephrine (NE) content in the dorsal hippocampus (HPC) and combined prelimbic and infralimbic cortices (PFC) depending on sex and hormonal status. A) OPFR-treated males demonstrated a reduction in HPC DA content in contrast to control males. B) OPFR-treated females in low steroid states displayed a trending decrease in PFC DA content in contrast to control-treated low-steroid females. Moreover, OPFR-treated females in high steroid states illustrated an increase in PFC DA content in contrast to control-treated high-steroid females. C) No pairwise differences were observed on HPC NE content. D) On the examination of PFC NE content, OPFR-treated females in high steroid states displayed a trending reduction in contrast to control-treated high females. Data are represented as mean ± SEM. * = *p* < .05; t = *p* < .10.

#### 3.7.2 Norepinephrine

On the examination of NE content in the dorsal hippocampus (Fig. 7C), we did not reveal an effect of treatment, hormonal status, or an interaction between treatment and hormonal status. On the analysis of NE content in the combined prelimbic and infralimbic cortices (Fig. 7D), we did not detect an effect of treatment, hormonal status, or an interaction between treatment and hormonal status. However, pairwise comparisons did reveal a trending reduction in NE content in OPFR-treated high-steroid females in contrast to control-treated high-steroid females (*p* = .0830).

### 3.8. RNA Sequencing of the Hippocampus (HPC)

#### 3.8.1 Total Gene Expression was Lower in OPFR-treated Mice

Total counts of gene transcripts were significantly lower in OPFR-exposed subjects [Oil: 27,694,954 ± 4,553,760; OPFR: 23,383,514 ± 3,866,509; *t*(27.28) = 2.795, *p* < .01., *d* = 1.02]. There was no interaction effect of sex or hormone status on total gene expression.

#### 3.8.2 OPFR Treatment Caused Differential Gene Expression (DGE) Profiles in the HPC

##### 3.8.2.1 Treatment-wide Differentially Gene Expression

We approached differential gene expression analysis by comparing the effect of OPFR treatment between all hormone state groups combined and between all hormone state groups separately. Of 12,113 genes, 13 had lower levels of transcript in OPFR-treated subjects and 108 had higher levels compared to Oil treated mice. A volcano plot of gene expression (Fig. 8A) and a selection of DEGs with the highest log 2-fold change and lowest eFDR are highlighted, separated by up-and downregulation. Values of log2FC and eFDR for top genes are reported (Table 1). Gene Ontology (GO) terms are also reported (Fig. 8B), displaying functional modules associated with clusters of highly differentially expressed genes.

**Fig. 8.**
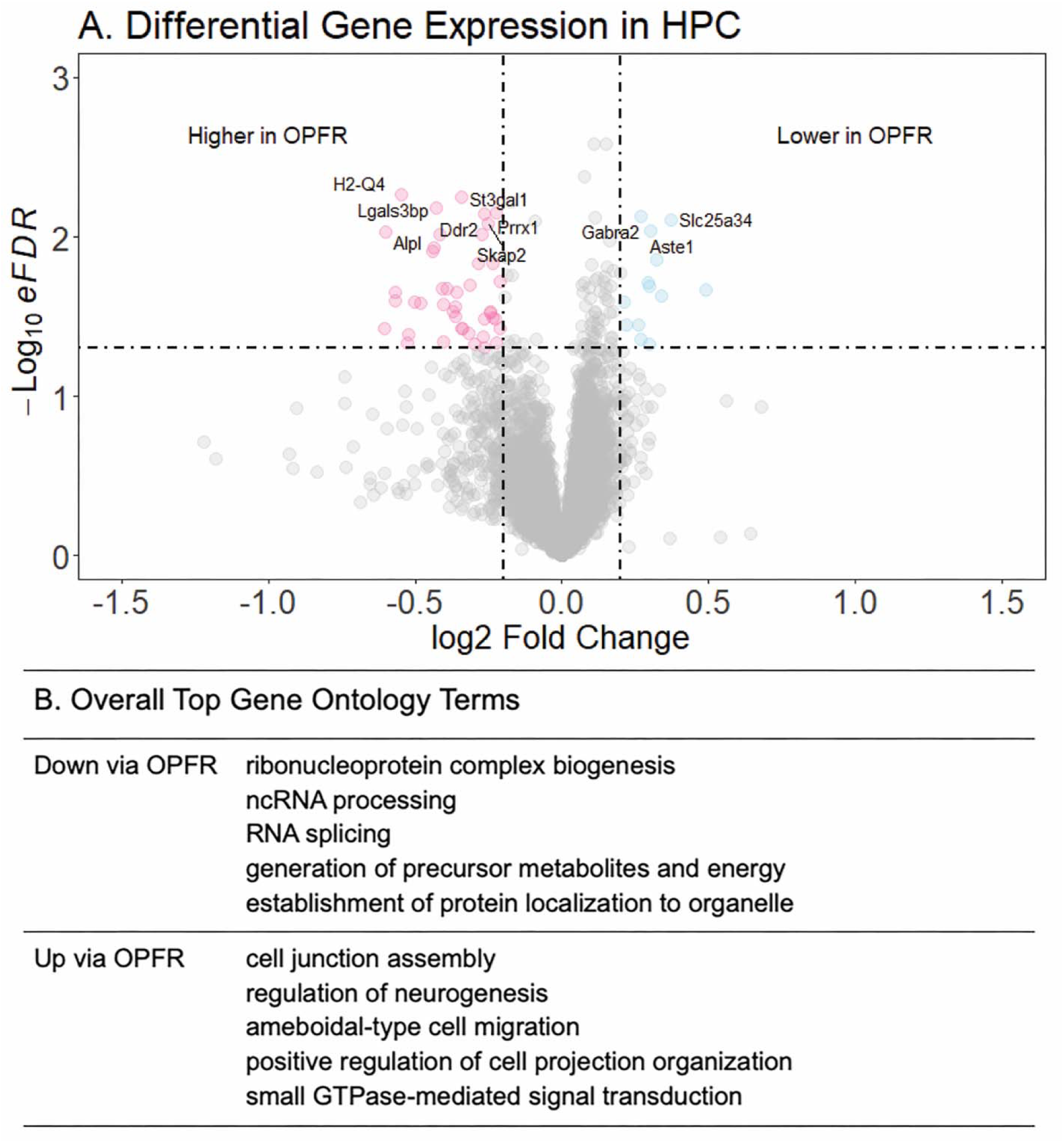
Results of Differential gene expression (DEGs) comparing all control and OPFR-exposed subjects together. Genes were determined to be differentially expressed if the log2 Fold Change was >= |0.2| and significant (eFDR) < 0.05. Functional modules identified by Gene Ontology enrichment analysis (GO terms) are below.

**Table 1.**
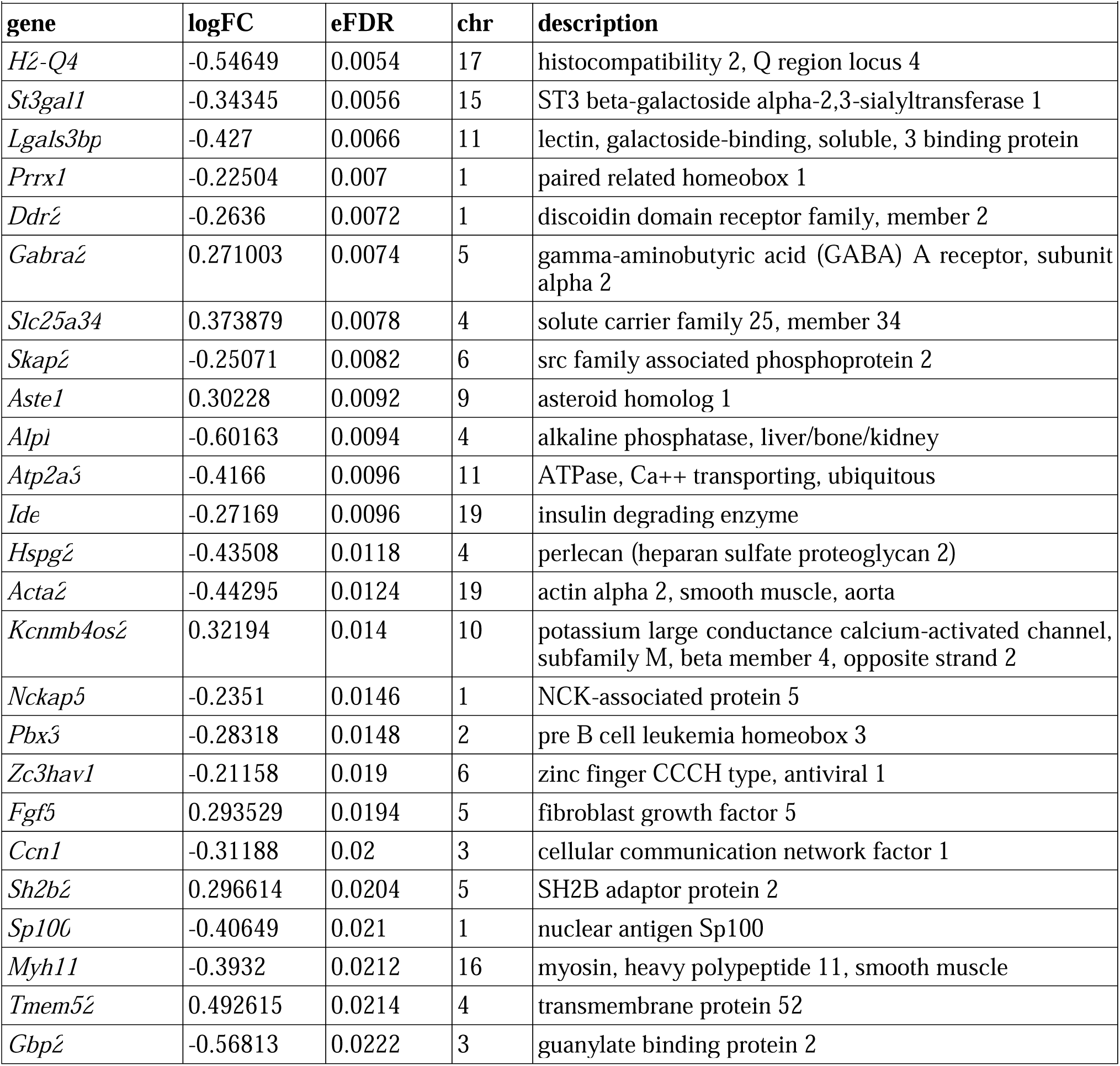
Top differentially expressed genes by OPFR treatment between all hormone state groups combined. Summary of top highly differentially expressed genes when comparing all control and OPFR-exposed subjects together, sorted by magnitude of significance. log2FC: log2 fold change in expression between groups. Positive values indicate lower levels of transcript in OPFR-exposed subjects; negative values indicate higher levels in OPFR-exposed subjects. eFDR: enhanced false discovery rate, or permuted p-value. chr: chromosome number location of gene.

##### 3.8.2.2 OPFR-induced Differential Gene Expression is Distinct Between Hormone States

Volcano plots of gene expression and a selection of DEGs with the highest log 2-fold change and lowest eFDR are highlighted, separated by up- and downregulation (Fig. 9). Values of log2FC and eFDR for top genes across all hormone status groups are reported (Supplemental Table 1). Gene Ontology (GO) terms are also reported (Fig. 9), displaying functional modules associated with clusters of highly differentially expressed genes. Within males, 70 genes were lower in transcript in OPFR-treated subjects and 70 were higher (Fig. 9A). Within low-steroid females, 142 genes were downregulated in OPFR-exposed and 79 were upregulated (Fig. 9B). Within high-steroid females, 117 genes were downregulated in OPFR-exposed and 90 were upregulated (Fig. 9C).

**Fig. 9.**
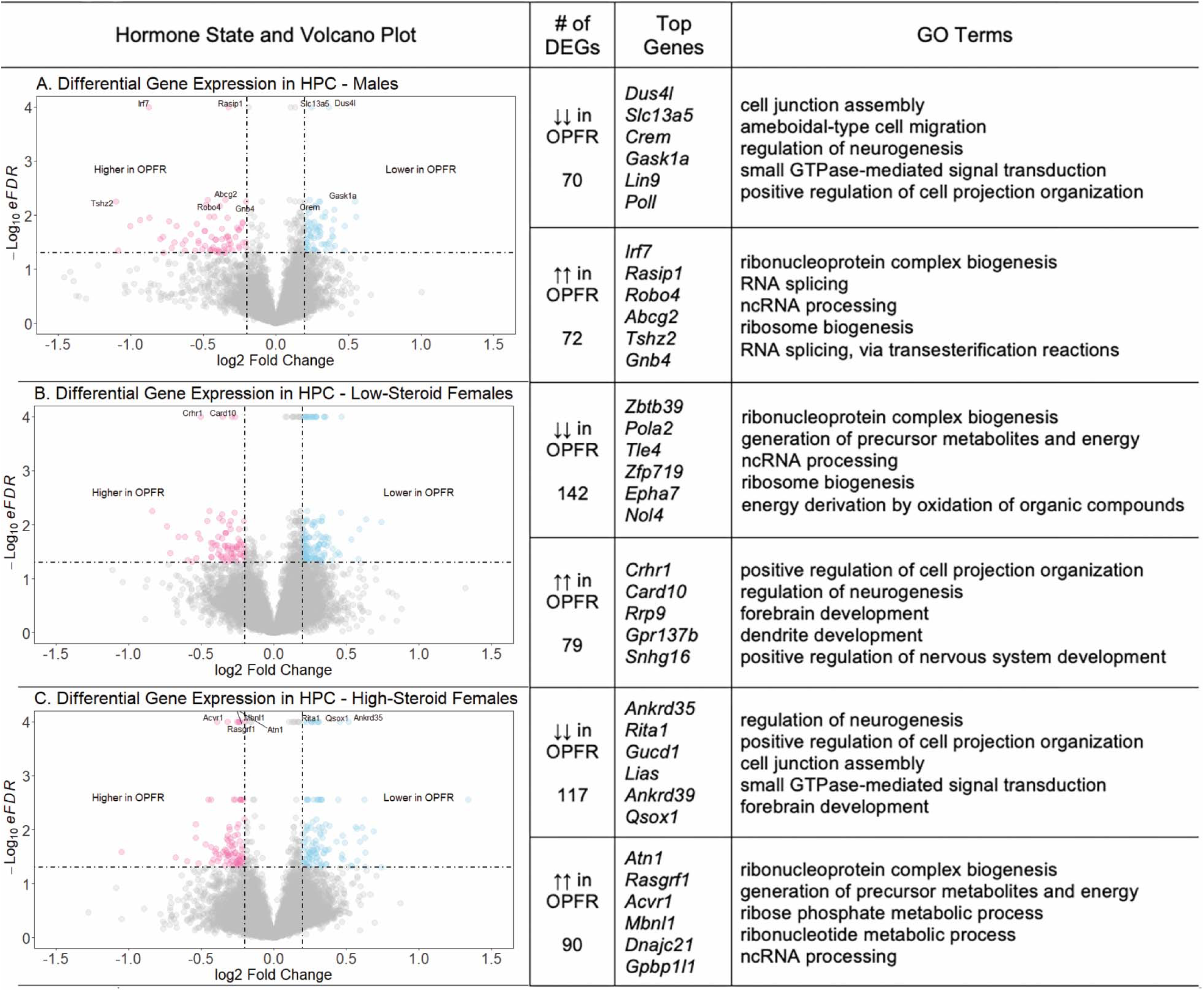
Results of differential gene expression (DEGs) when comparing A. Males, B. Low steroid state females, and C. high steroid state females. Genes were determined to be differentially expressed if the log2 Fold Change was >= |0.2| and the significant (eFDR) < 0.05. Top genes were identified as the greatest absolute value log2FC with an eFDR < 0.05. Functional modules identified by Gene Ontology enrichment analysis (GO terms) are on the right.

#### 3.8.3 Frequency of GO Term Genes

As each GO term is a collection of genes related to a certain biological function, different GO terms may contain the same gene. We compiled the genes that occur most frequently in the top 100 up- and downregulated GO terms for each hormone state group to give a better sense of genes that are involved in multiple functions altered by OPFR exposure (Table 2). While these genes may not be differentially expressed between Oil and OPFR groups, their frequent occurrence in differentially expressed GO terms is notable.

**Table 2.**
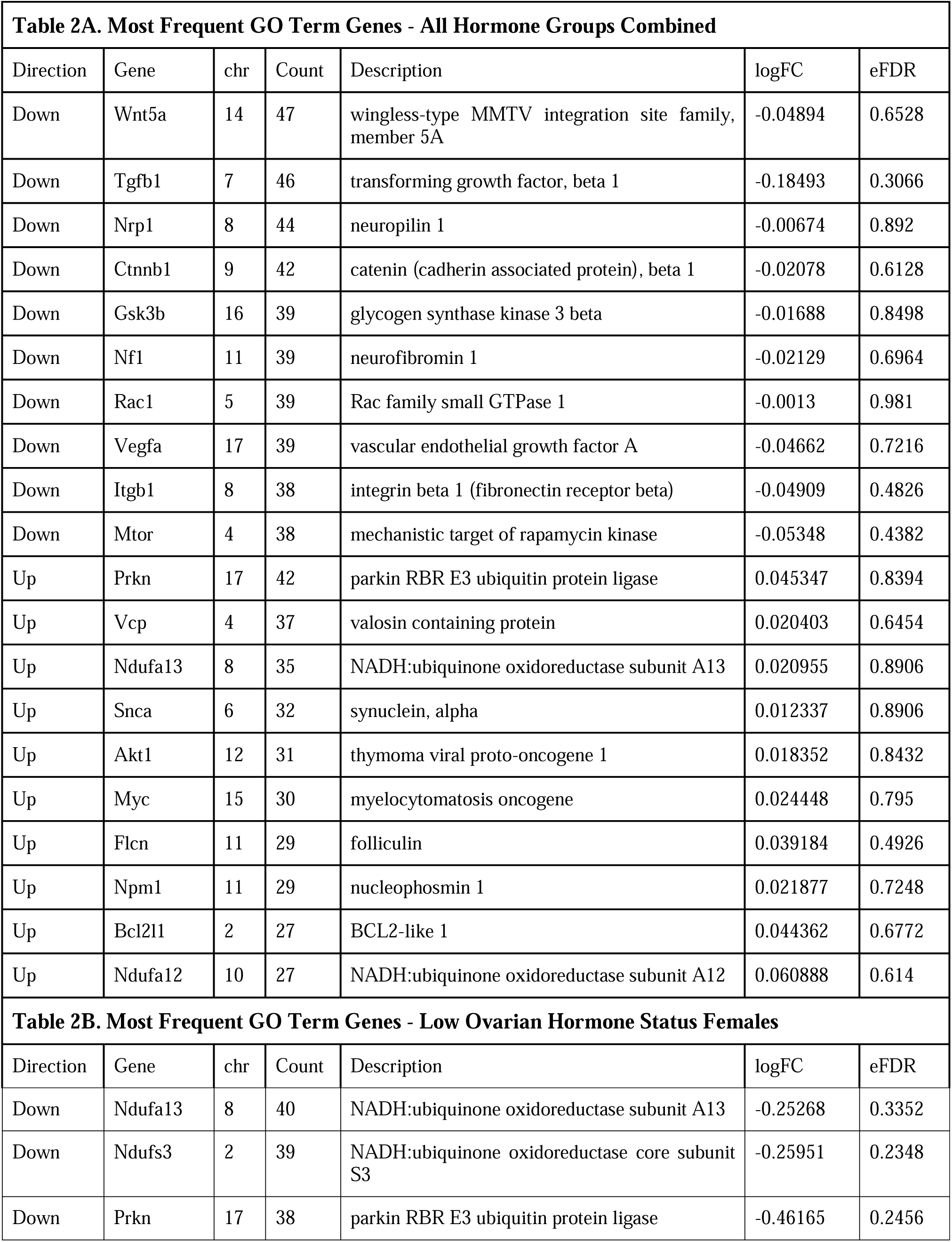
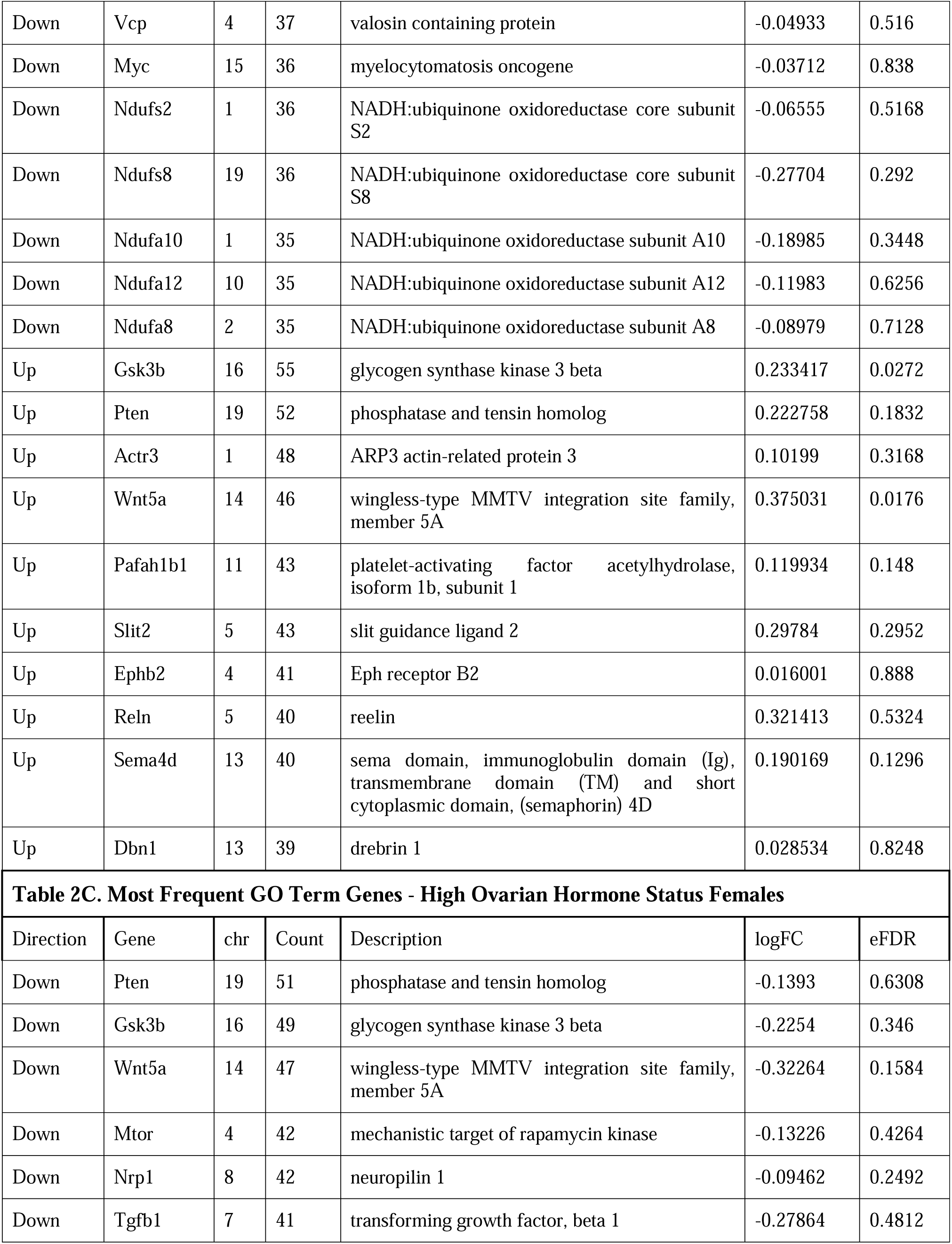
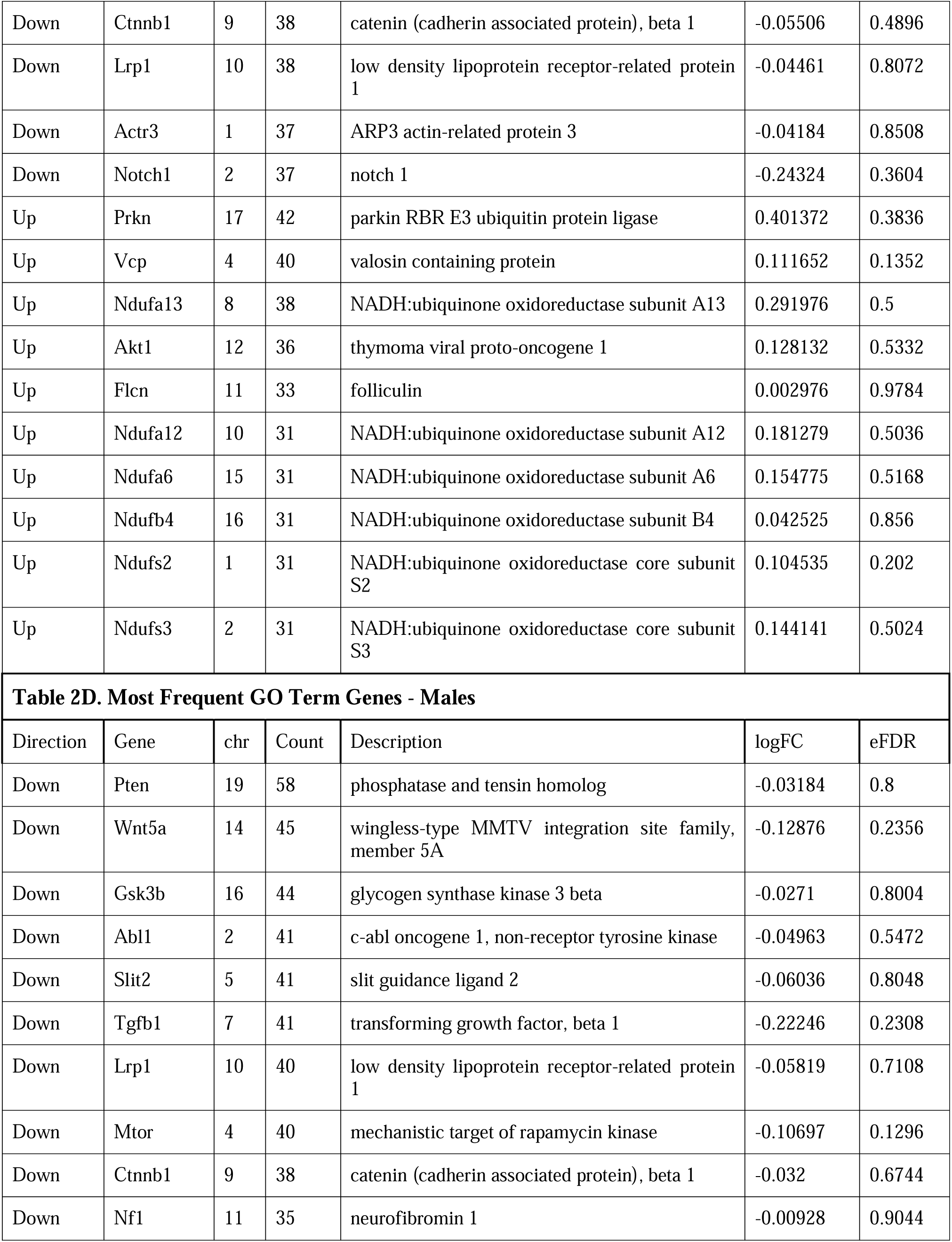
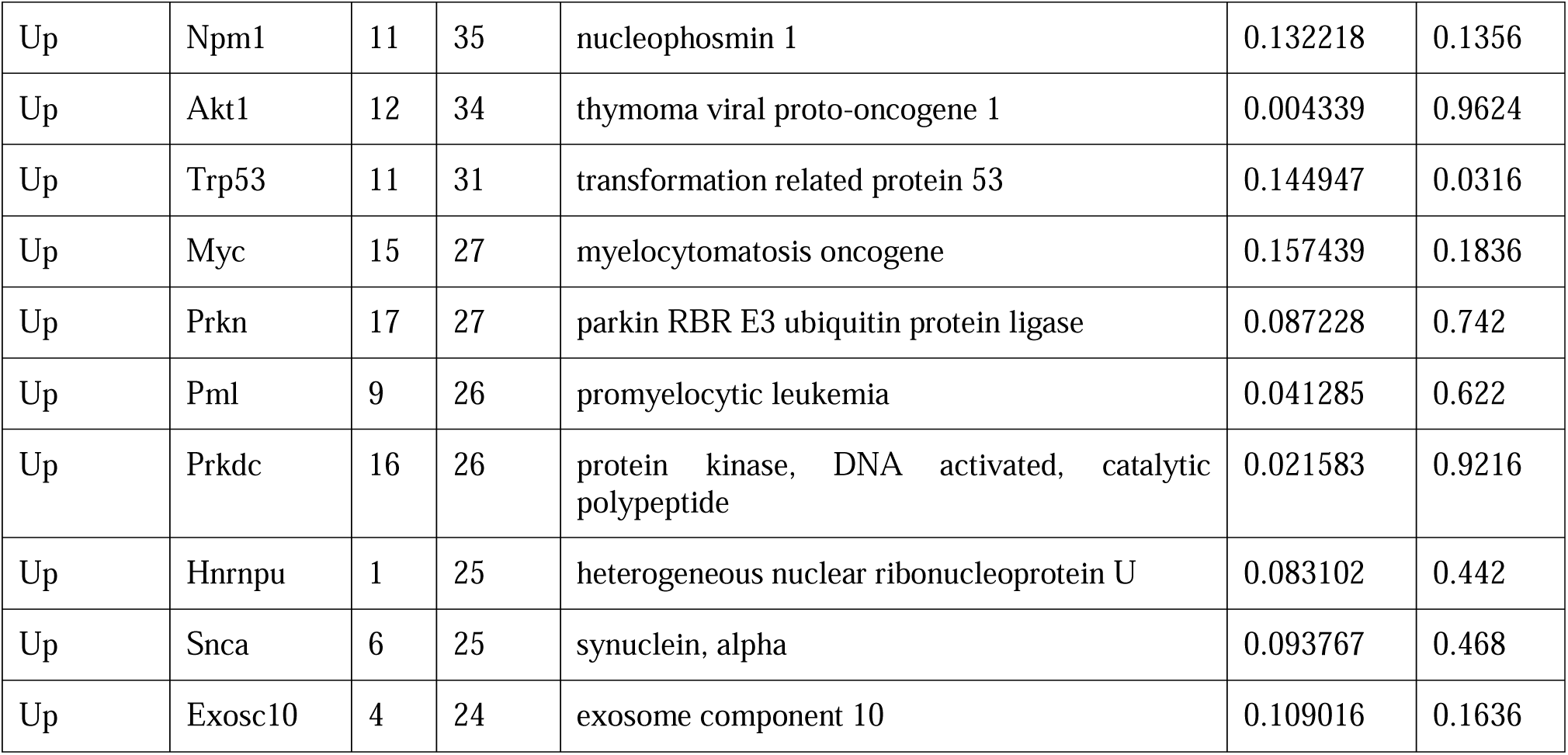
Compilation of genes most frequently associated with the top 100 up- and downregulated GO terms. Direction: Down indicates that the GO terms the gene was involved in was downregulated, Up indicates that the GO terms the gene was involved in was upregulated. Count: The number of GO terms the gene was involved in. log2FC: log2 fold change in expression between groups. Positive values indicate lower levels of transcript in OPFR-exposed subjects; negative values indicate higher levels in OPFR-exposed subjects. eFDR: enhanced false discovery rate, or permuted p-value. chr: chromosome number location of gene.

#### 3.8.4 DGE Overlap and Contrast

We also performed an overlap and contrast analysis on DEGs, compiling genes that were increased or decreased in expression in multiple hormone states (Overlap) (Table 3A), and genes that showed increased expression in one hormone state but decreased expression in another (Contrast) (Table 3B).

**Table 3.**
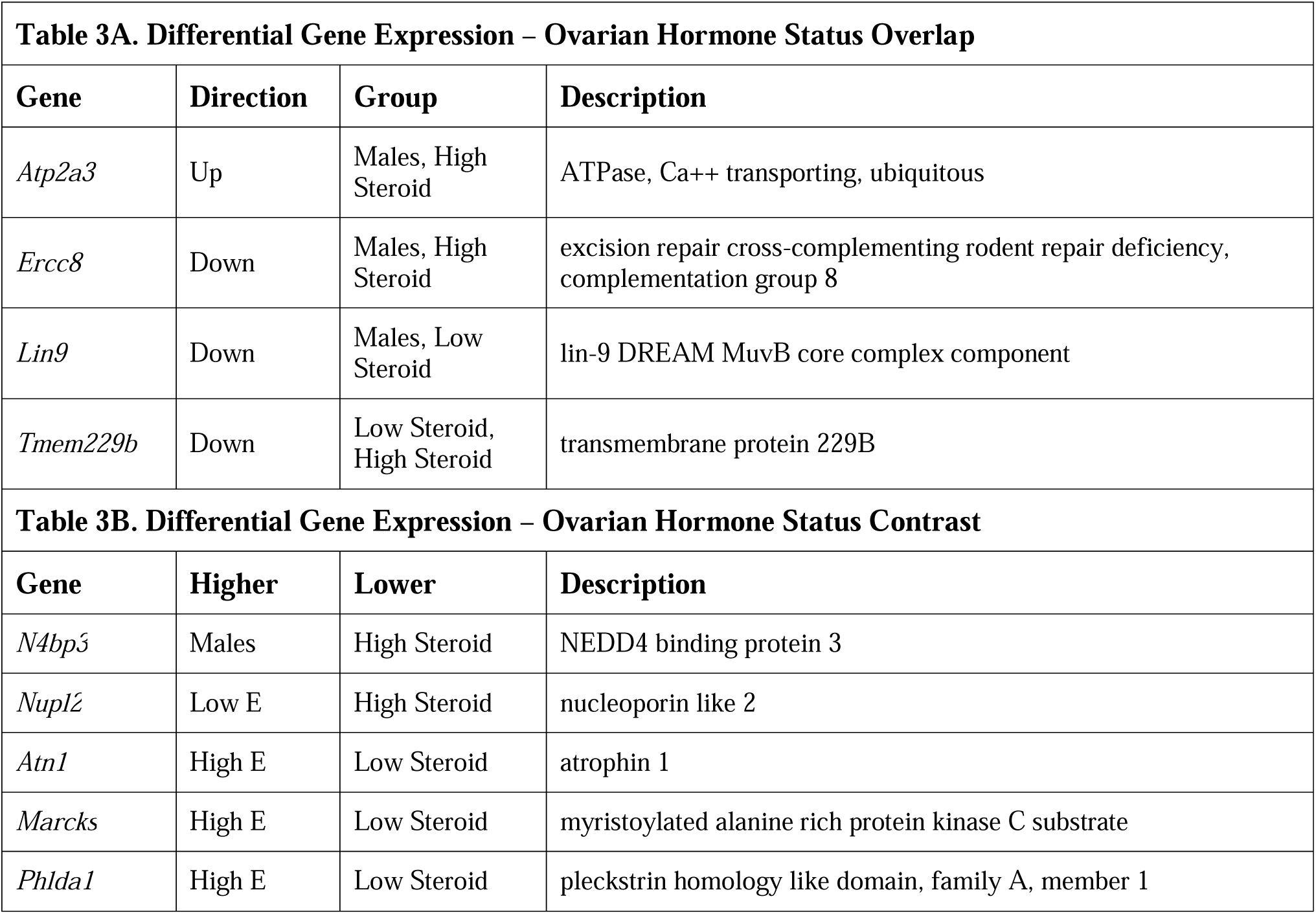
Overlap and Contrast of genes differentially expressed due to OPFR exposure between Ovarian Hormone Status groups. In A. Overlap, direction indicates if genes were up- or downregulated, and group identifies which Hormone Status groups share those overlaps. In B. Contrast, the list is first organized by which group contrasting genes were upregulated in OPFR-exposed subjects, then which groups they were downregulated in.

#### 3.8.5 WGCNA Highlights Distinct Functional Modules Overexpressed in OPFR-treated Mice

Similar to DGE, we approached WGCNA by comparing the effect of OPFR treatment between all ovarian hormone status groups combined and within all treatment and hormone groups separately. Summaries to all WGCNA modules are identified (Fig. 10). Modules differentially expressed between groups are highlighted (Table 4), including the GO terms most associated with genes in the module. Genes with high module membership (MM - genes that are most highly correlated with other genes in the module) are also listed. In whole-treatment analysis, the salmon module (Table 4A) was significantly upregulated in OPFR subjects (b = 0.151 ± 0.065; n = 29, *p* < .05). When comparing all groups, the pink module (Table 4B) was higher in OPFR-treated males (b = 0.193 ± 0.087; n = 29, *p* < .05) and the black module (Table 4C) was lower in control-treated high ovarian hormone status females (b = -0.187 ± 0.088; n = 29, *p* < .05).

**Fig. 10.**
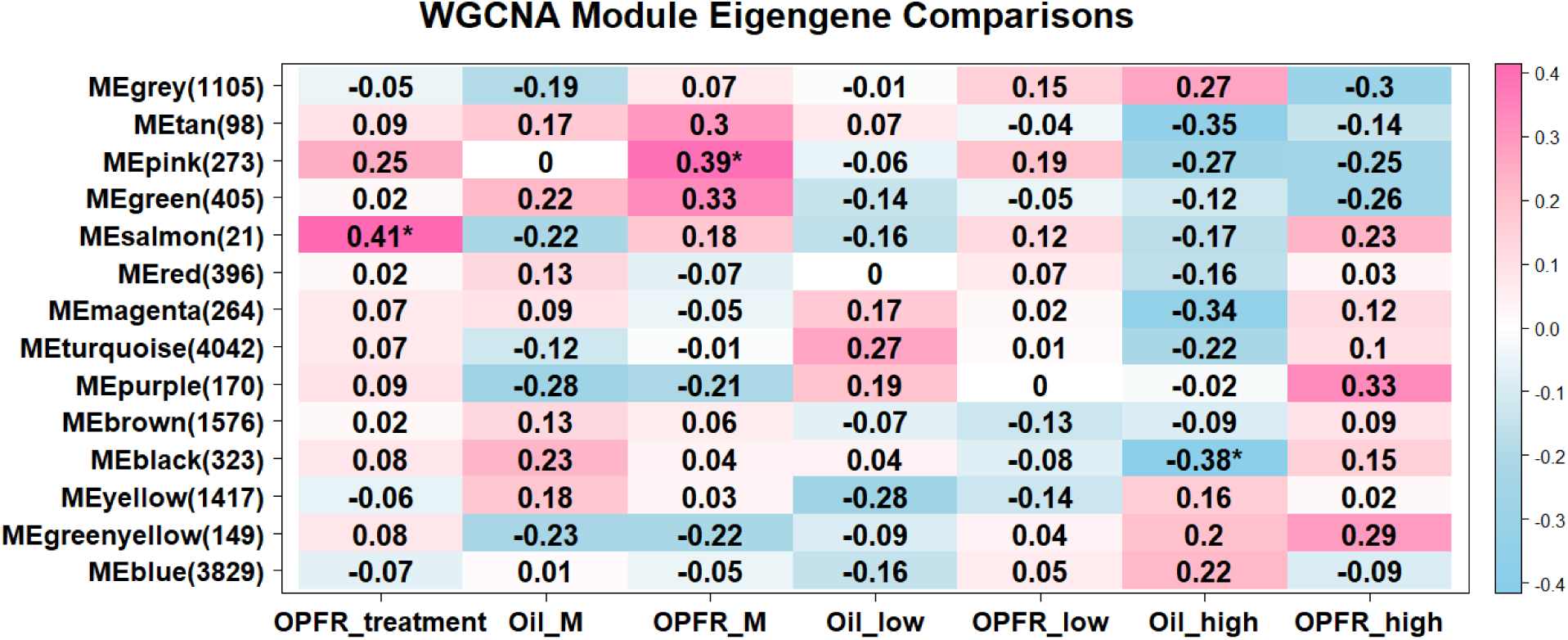
Results of weighted gene coexpression network analysis (WGCNA) comparing all control-treated and OPFR-treated subjects together (OPFR_treatment) and each treatment x ovarian hormone status group. On the left are the modules identified, next to the number of genes in those modules. In the OPFR_treatment column, color of boxes represents whether a module is upregulated (pink with a positive value) or downregulated (blue with a negative value) in OPFR-treated subjects, with the darkness of the shade reflecting magnitude. Significantly different modules are marked with an asterisk (*). In the treatment * status columns, same as above, except color and shade signifies the difference in module expression in the group named in the column compared to all other treatment * status groups.

**Table 4.**
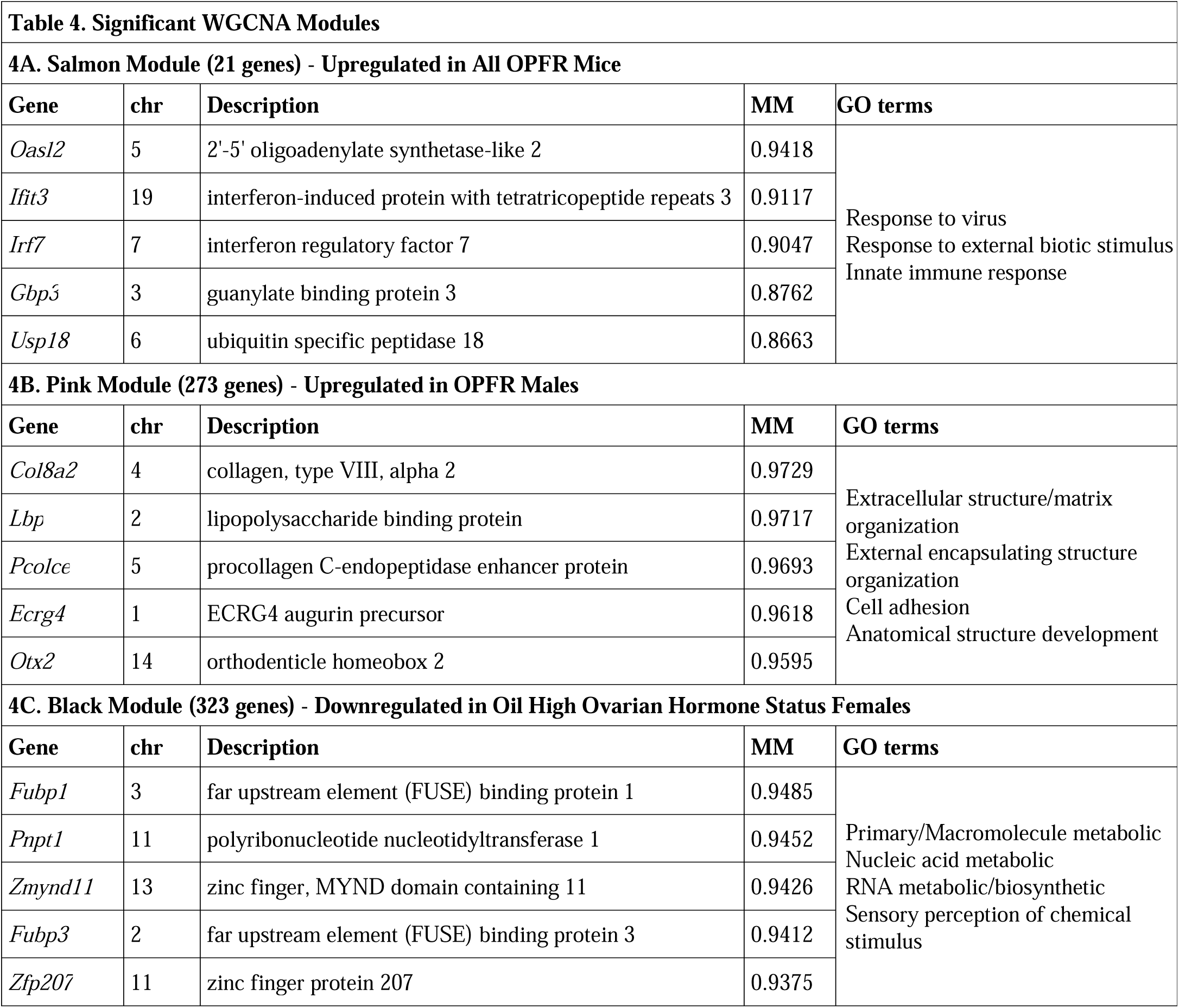
Significantly differentially expressed WGCNA-derived modules across different comparisons. A. The salmon module was upregulated in all OPFR-exposed groups. B. The pink module was upregulated in OPFR-exposed males compared to all other groups. C. The black module was downregulated in oil-treated high ovarian hormone status females compared to all other groups. Gene Ontology (GO) terms associated with differentially expressed modules is on the right, with hub genes with highest module membership (MM) and the chromosome location (chr) of those genes on the left.

## 4. Discussion

Perinatal exposure to OPFRs produced long-lasting, sex- and hormone-dependent alterations in anxiety-like behavior, memory, neurotransmitter content, and gene expression profiles in male and female mice. Across several behavioral paradigms such as the Y-maze, SOR, NOR, and the Barnes maze we observed behavioral alterations dependent on sex and ovarian hormone status. These findings provide evidence that perinatal exposure to the OPFR mixture, tris(1,3-dichloro-2-propyl) phosphate (TDCPP), triphenyl phosphate (TPP), and tricresyl phosphate (TCP) can modify neural and behavioral trajectories in a sex-specific and hormone-dependent manner. Moreover, differential dopaminergic and noradrenergic responses in the hippocampus and prefrontal cortex suggest that OPFRs exert enduring effects on monoaminergic signaling pathways that may mediate the observed behavioral outcomes. Perinatal OPFR exposure also causes a variety of differential gene expression with distinct profiles between ovarian hormone (steroid) status groups. Collectively, these results highlight the neurotoxic effects of OPFRs during development.

### 4.1 Anxiety

Prior findings demonstrate that perinatal exposure to organophosphate flame retardants alters anxiety phenotypes in a sex-specific manner [24, 30, 31, 65, 66]. Several of these studies utilized FireMaster® 550, a flame retardant mixture that contains TPP as used in our study. Previous work in prairie voles (*Microtus ochrogaster*) that were perinatal exposed to FireMaster® 550 (500, 1000, 2000 μg/20 μl; via sc injection) from GD 0 to PND 21 were tested as adults (after PND 80) on the OFT [66]. The 500 and 2000 μg FireMaster® 550 females entered the center less often than controls suggesting an anxiogenic phenotype. The 1000 μg females also made less center entries but, did not reach statistical significance [66]. Baldwin and colleagues [24] also examined the effects of developmental exposure to Firemaster® 550. Dams were administered from GD 9 to 18 and authors reported sex-dependent alterations in anxiety-like behavior in adult Wistar rats. Adult males displayed heightened anxiety in the EPM, characterized by reduced time and entries in the open arms, whereas females, tested in estrus, exhibited increased anxiety-like behavior specifically in the light–dark box, demonstrated by delayed transitions into the light side of the chamber. Similarly, Patisaul et al. [30] identified sexually dimorphic outcomes following perinatal exposure to Firemaster® 550 from GD 8 through weaning. In adulthood, females, tested in estrus, exhibited greater anxiety-like behavior, reflected by reduced open arm exploration on the ZM at the 100-μg doses. In contrast, males exposed to the lower 100-μg dose showed increased open arm ZM activity, indicative of an anxiolytic-like effect. Moreover, Witchey et al. [65] exposed Wistar dams to Firemaster® 550 (2000 μg/day) 72 h after pairing to PND 21. Firemaster® 550 exposed females, tested in diestrus, illustrated a decreased latency to enter the center of the OFT suggesting lower anxiety, while Firemaster® 550 exposed males exhibited no differences. Our previous work [31] demonstrated that maternal exposure to our same OPFR mixture (1mg/kg/day) used here elicited sex-specific effects on anxiety-like behavior in adult offspring. OPFR-treated male mice exhibited an anxiolytic phenotype in the elevated plus maze (EPM) as evidenced by an increased number of open arm entries, whereas female behavior was unaffected. Here, our findings suggest that developmental exposure to our OPFR mixture modulates anxiety-like behavior in females in a manner dependent on ovarian hormone status. Although we did not observe any differences in males on the OFT, OPFR-treated females tested in high steroid states (proestrus/estrus) exhibited reduced time in the center of the open field suggesting increased anxiety-like behavior. Conversely, OPFR-treated females tested in low steroid states (metestrus/diestrus) illustrated an inverse anxiety pattern, revealing a trend in increased 10cm center exploration indicative of an anxiolytic phenotype. Interestingly, our finding in the low ovarian status females are most similar to Witchey et al.[65] who found an OFT anxiolytic phenotype from perinatal Firemaster® 550 exposed female Wistar rats tested in diestrus. Moreover, the anxiogenic effects reported by Baldwin et al. [24] and Patisaul et al. [30] in females tested during the estrous phase parallel our observations of increased anxiety-like behavior in high steroid females. Collectively, these findings suggest a hormone-dependent OPFR-induced reversal in anxiety-like behavior in females. This bidirectional pattern indicates that perinatal OPFR treatment alters the developmental organization of neural circuits that later respond to ovarian hormones. Such disruption may impair how the adult brain integrates endocrine signals to regulate anxiety-like behavior. One plausible mechanism is that perinatal OPFR interferes with estrogen receptor signaling, neurosteroid synthesis, or related transcriptional pathways during critical periods of neural differentiation, thereby reprogramming the sensitivity of hormone-responsive brain regions. In addition, perinatal OPFRs may induce long-lasting alterations in neurotransmitter systems such as noradrenergic or dopaminergic networks that interact with ovarian hormones across the estrous cycle. Our results suggest that perinatal OPFR exposure developmentally reconfigures the neuroendocrine system, resulting in persistent changes producing opposing patterns of anxiety-like behavior in adulthood depending on ovarian hormone status.

### 4.2 Memory

#### 4.2.1 Y-Maze

We also utilized the Y-Maze to test short-term reference memory for spatial navigation. The Y-Maze evaluates spatial navigation by measuring a rodent’s ability to recognize and preferentially explore an unknown arm based on spatial cues after a delay period [67]. Our previous work supports the evidence that OPFRs disrupt spatial navigation memory. In our prior work [34], perinatal exposure to the same OPFR mixture and dose used in the current study (1mg/kg/day; GD7 to PND14; TPP, TDCPP, and TCP) produced lasting impairments in spatial memory in adulthood. OPFR-treated female offspring maintained on a high-fat diet exhibited a significant reduction in the percentage of time spent exploring the unknown arm of the Y-maze relative to controls, consistent with deficits in spatial memory performance. We also previously found that perinatal OPFR-treated male offspring did not display measurable alterations in spatial memory [34]. Our findings in the current study align with our prior work in the Y-Maze. OPFR-treated high-steroid females exhibited a reduction in time spent exploring the unknown arm of the Y-Maze relative to control high females suggesting compromised spatial memory performance. Similarly, OPFR-treated males were unaffected in the Y-Maze of our current study. However, because Yasrebi et al. [34] did not differentiate estrous cycle stages, direct comparison across studies remains limited, as here we also uncovered no differences in our low ovarian hormone status females. It is possible that the females in Yasrebi et al. [34] were predominantly in high hormone stages, which could account for the observed similarities in behavioral outcomes. A relevant study independent of our own research administered several concentrations of TPP (0.5, 5, and 50 mg/kg) orally from PND 10 to 70 in wild-type (C57BL/6) [36]. This research utilized a modified Y-Maze paradigm that evaluated correct spontaneous alternation behavior. Although the sex of the mice was not reported, Zhong et al. [36] revealed TPP treatment lowers the percentage of spontaneous alteration behavior at the two highest doses indicative of spatial navigation and reference memory deficits. Differences in findings between Zhong and colleges [36] and our own work may be attributed to variations in experimental design, as the authors examined only TPP, whereas our study used a mixture of OPFRs (TPP, TDCPP, and TCP) that may exert broader or interacting neurotoxic effects. Also, their use of an alternative Y-Maze paradigm and a postnatal only exposure window likely engaged different cognitive domains and developmental periods than those assessed in our perinatal exposure model.

#### 4.2.2 Spatial Object Recognition (SOR)

In addition to the Y-Maze, we also employed SOR to further evaluate perinatal OPFR treatment on hippocampal-dependent spatial memory and its modulation by sex and hormonal status. SOR examines a rodent’s ability to detect changes in the spatial arrangement of known objects [68]. It evaluates how well a rodent encodes, retains, and retrieves information about object orientation. In the current study, OPFR-treated males demonstrated trending and significant reductions in both time spent and number of interaction bouts with the displaced object, respectively. Moreover, these males also displayed a trending decrease in the percentage of displaced object bouts. Although we observed no differences in females in low-steroid states, females in high-steroid states demonstrated a significant decrease in the percentage of displaced interaction object bouts relative to controls. To the best of our knowledge, no previous studies have evaluated the effects of perinatal exposure to the specific OPFRs used in our mixture on spatial object recognition performance. Nevertheless, our findings indicate that there are both sex- and hormone-dependent impairments in spatial object orientation memory. Considering the outcomes from both our Y-maze and SOR tests, which are spatial assays conducted under low stress conditions, our memory deficits in high females suggest that elevated ovarian hormone levels may exacerbate the cognitive effects of OPFR exposure. This effect is likely driven by interactions between endocrine signaling and hippocampal-dependent mechanisms that support spatial information processing.

#### 4.2.3 Novel Object Recognition (NOR)

In addition to spatial memory assessments, we also evaluated recognition memory using the NOR test. The NOR test measures recognition memory by evaluating a rodent’s ability to distinguish between familiar and unfamiliar objects based on prior experience. Rodents naturally exhibit a preference for novelty; therefore, when they recognize the familiar object, they preferentially allocate more time exploring the novel one [69]. As our and other’s [48–50] paradigm applies repeated habituation to minimize reliance on spatial cues, we consider this test to be largely hippocampal-independent. The aforementioned work by Zhong and colleagues [36] examining oral TPP (0.5, 5, and 50 mg/kg) from PND 10 to 70 in wild type mice also tested mice on NOR. Again, the sex was not reported; however, both the duration of exploration of the novel object and the discrimination index declined at the two highest TPP exposure levels relative to controls [36]. In addition, Hawkey and colleagues [33] evaluated 7 week old Sprague–Dawley rats that were perinatally exposed to TPP (16 or 32 mg/kg; throughout pregnancy up to 8–11 days into the neonatal stage) on the NOR test. Hawkey et al. [33] revealed that for the high TPP dose there was significantly less novel object preference compared to controls suggesting an impaired recognition memory. The authors included male and female subjects; however, the interaction between sex and treatment did not reach significance. The aforementioned research [66] that perinatally exposed Prairie voles to FireMaster® 550 (500, 1000, 2000 μg/20 μl) from GD 0 to PND 21, tested the voles on the NOR. Of the FireMaster® 550 treated females, recognition was impaired in the 500 and 2000 μg females in both tests, a 30-minute short term and 24-hour long-term memory test. When testing males, recognition memory was impaired in all but the 500 μg dose males in the short-term memory test. No differences were observed in NOR long term memory for males [66]. In contrast to these previous studies, our findings did not reveal robust deficits in recognition memory following perinatal exposure to our OPFR mixture. However, we observed a trend toward increased novel object interaction bouts in OPFR-treated high ovarian hormone status females relative to controls, which may indicate an improvement in memory. Although, as this is only a trending effect, it may reflect subtle hormone-dependent alterations in exploratory drive or novelty-seeking behavior in this test rather than a true cognitive alteration.

#### 4.2.4 Barnes Maze

Deficits in spatial learning and memory following developmental FR exposure to the polybrominated diphenyl ethers have been reported across rodent models. Sun et al. [70] demonstrated that developmental exposure (GD 1 to 21) to the polybrominated diphenyl ether, BDE-209, in Sprague–Dawley rats impaired spatial learning in the Morris water maze, which also measures hippocampal-dependent spatial memory like the Barnes Maze, but is more stressful [71]. PND 28 offspring displayed prolonged escape latencies from exposure doses BDE-209 10 and 20 mg/kg/day. However, because the authors did not specify the sex of the tested offspring, interpretation of these findings in relation to our sex- and hormone-dependent results remains limited. There is limited work that investigates perinatal OPFR exposure on studies similar to the Barnes maze specifically. Our Barnes maze results highlight the complexity of OPFR-induced spatial memory alterations. In the short-term Barnes maze, OPFR-treated males spent less time in the target quadrant suggesting a memory deficit. Interestingly, females in high ovarian hormone states showed a decreased escape latency suggesting an improvement in this type of memory. As no prior studies have investigated the effects of perinatal OPFR exposure on Barnes maze performance with respect to sex or hormonal status, the underlying mechanisms driving these differences remain unclear. Females in high ovarian hormone states may have had improved short-term Barnes maze results because elevated steroid levels enhance both hippocampal plasticity and stress regulation, two systems critical for spatial memory under moderately stressful conditions. Notably, the estrous cycle is characterized by dynamic changes in dendritic spine density within the hippocampal CA1 region. Spine density and synaptic number reach their highest levels during the proestrus phase or following exposure to exogenous estradiol [72]. At the same time, females exhibit cognitive resilience to chronic stressors (repeated Barnes maze training) [73]. Thus, moderate stress often enhances female performance in spatial memory tests. Because the Barnes maze relies on aversive motivation (open, elevated space), females with high circulating steroids may experience reduced stress reactivity, allowing for more effective spatial cognitive engagement. Thus, females in high ovarian hormone states likely have improved Barnes maze performance due to a combined effect of estrogen enhanced hippocampal function and attenuated stress-induced cognitive interference.

Evaluating the long-term Barnes maze in the current study, females in both ovarian hormone groups revealed no differences. OPFR-treated males had a reduction in escape hole entries in contrast to their control counterparts suggesting long-term memory disfunction. One plausible explanation behind only males exhibiting deficits during the long-term Barnes maze may relate to sex-specific differences in how OPFRs affect long-term hippocampal-dependent memory consolidation and retrieval processes over time. Long-term memory relies critically on synaptic plasticity, dendritic spine maintenance, and long-term potentiation within the hippocampus; mechanisms that are sensitive to endocrine disruption [74]. Males may be particularly susceptible to these disruptions due to lower baseline estrogenic neuroprotection. In contrast, females, may experience protection granted by 17β-estradiol mediated enhancement of hippocampal plasticity and spatial memory retention [75]. Consequently, the absence of such hormonal buffering in males could explain why OPFR impairments are present during long-term memory testing in the Barnes maze.

### 4.3 Neurotransmitter Content

Our neurochemical findings provide additional insight into potential mechanisms underlying the observed behavioral changes. In males, perinatal OPFR treatment decreased DA levels in the HPC, whereas in females, HPC dopaminergic content remained unaltered in both groups of females. A reduction in HPC DA levels is associated with impaired long-term memory, consistent with our findings in the Barnes maze, where deficits in long-term memory performance were observed selectively in males. The hippocampus and ventral tegmental area (VTA) form a functional loop in which hippocampal detection of novel information triggers dopaminergic activation in the VTA. Dopamine released back into the HPC enhances long-term potentiation and learning, thereby regulating how new information is encoded into long-term memory (reviewed in [76]). Together, these results suggest that perinatal OPFR treatment may disrupt HPC–VTA dopaminergic signaling in a sex-dependent manner, leading to long-term memory impairments in males. In the PFC, OPFR-treated females in high steroid states demonstrated increased PFC dopaminergic content but had several differences in behavioral endpoints. Dopamine in the PFC follows an inverted-U relationship with cognition, where optimal levels support working memory, but excessive DA impairs these functions (reviewed in [77]). In OPFR-treated females in high steroid states, increases in PFC DA may have elevated DA signaling beyond the optimal range, contributing to the poorer performance observed in the Y-Maze and SOR, which rely on flexible spatial updating and attention to spatial detail. Although these tasks typically rely on the HPC, PFC involvement may impact performance, especially in the integration of contextual information [78, 79].

### 4.4 Perinatal OPFR Treatment Causes a Variety of Gene Expression Differences

Transcript expression of *gabra2*, gamma-aminobutyric acid (GABA) A receptor subunit alpha 2, was lower in OPFR-treated subjects regardless of hormone state. GABA is the most common inhibitory neurotransmitter, with widespread function throughout the brain. While EDC-induced up- and downregulated GABA-related expression has occasionally been noted in other regions, such as the amygdala, and has been associated with ADHD-like behavioral traits [80, 81], the causal relationship remains unclear. However, allelic *gabra2* expression differences in humans have frequently been associated with drug and alcohol use likelihood [82, 83], and these behavioral traits are largely correlated with anxiety and cognitive impairment. While *gabra2* was the only GABA related gene to be differentially expressed in OPFR-treated subjects, others have shown that similar OPFR mixtures in neuronal culture models alter *gabra1* transcript [84], demonstrating that the effects of OPFRs can be selective on specific subunit expression. Nevertheless, while *gabra2a* transcript was lower in OPFR-treated animals, many of the cognitive and mood related behaviors were only altered in a hormone-state specific way, which we will discuss below.

Additionally, the top Gene Ontology terms upregulated by OPFR exposure were related to neurogenesis and the organization of cell migration, projection, cell junction assembly, and GTPase-modulated signal transduction. As the samples in the present study were from whole dorsal hippocampus, this could suggest significant effects of perinatal OPFR exposure on dentate gyrus (DG) function. Others have also shown that perinatal OPFR exposure causes changes in dentate gyrus volume in males and neurodevelopment markers in males and females [35, 36]. In males and females exposed perinatally to OPFRs, doublecortin expression, important for proper neurodevelopment, was decreased in the DG [35]. However, nestin, which is primarily expressed in neuronal progenitor cells, was downregulated in males but upregulated in females [35]. The present data provide additional support for these findings. In our OPFR-treated males, the WGCNA pink module was upregulated. The pink module contains genes involved with extracellular structure and matrix development, suggesting a possible mechanistic difference in organizational gene expression and activity. More broadly, alongside an overall downregulation of gene expression, lower cellular energy production-related gene transcription, and potential upregulation of inhibitory neurons (via GABAergic gene upregulation), increased neurogenesis processes may reflect a compensatory mechanism of decreased functional excitability in the OPFR-exposed developing hippocampus.

In some studies, OPFR exposure contributes to differences in gross metabolic function and body weights, sometimes reducing it [37], sometimes enhancing it when paired with a high fat diet [85], and sometimes not affecting it at all [86, 87]. However, metabolic effects are also dependent on time of exposure or sex and are paired with more subtle metabolic changes, such as changes to glucose and insulin tolerance. One of the most enriched Gene Ontology terms that was downregulated in OPFR hippocampal tissue was generation of precursor metabolites and energy, reflecting local changes to energy use. Furthermore, OPFR-treated subjects had increased transcript for *ide*, insulin degrading enzyme. Hippocampal insulin resistance is related to cognitive deficits in mouse models of type 2 diabetes and Alzheimer’s disease [88]. Hippocampal insulin resistance is driven, in part, by reduced insulin driven phosphorylation of Akt [89, 90]. In our OPFR-treated animals, *akt1* transcript was among the most frequently occurring genes in the top 100 most upregulated GO terms, especially in the males and high steroid state females. The *wnt* and *gsk3b* genes were also common among the most frequently occurring downregulated GO terms, again especially among males and high steroid state females. The *wnt/GSK3*β*/*β*-catenin* pathway is critical in early neurodevelopment and is sensitive to polycyclic aromatic hydrocarbon exposure [91]. The GSK3β pathway has also previously been associated with high fat diet induced hippocampal insulin resistance [92, 93]. Prior research has shown that perinatal OPFR exposure impairs insulin-induced glucose clearance in males, though this effect partially ameliorated the more drastic effect of a high fat diet exposure on insulin tolerance, especially in females [87]. Other genes, such as *Lgals3bp*, *H2-D1*, and *H2-K1* are also involved in the expression of insulin resistance in diabetes [94, 95]. Prior work from our lab has also demonstrated that OPFRs act on similar mechanisms as high fat diets and compound on the detrimental neuro-environment they cause [34, 96]. It would be worthwhile to investigate *Ide*, *Lgals3bp*, and other insulin-related genes in more tissues and contexts in future OPFR studies.

Many of the above genes associated with insulin resistance were included in the WGCNA salmon module, which was upregulated broadly in OPFR-exposed subjects. The major GO terms associated with genes in the salmon module were related to immune function. The salmon module contained a number of immune related genes, such as a variety of interferon protein and receptor coding transcripts such as *Irf7, Ifit3b, Ifitm3, Ifit3,* and *Ifit1*. The salmon module also contained genes related to extracellular matrix organization and cellular adhesion such as *H2-D1* and *H2-K1*, which are reactive to interferons [94]. Related histocompatibility genes, such as *H2-Q4*, though not within the salmon module, were also significantly higher in transcript level in OPFR-treated subjects. Recently, similar profiles of differential histocompatibility complex 2 gene expression were highlighted in a variety of brain areas in aged mice (18 months) compared to young mice (2-3 months) [97]. Galactose binding and processing related gene *Lgals3bp*, which is also involved in immune response pathways [95, 98], was also significantly upregulated in OPFR groups and a major component of the salmon module. Such differential expression aligns with other complications reported from a variety of EDC exposures that lead to immune and cognitive impairment [99, 100]. This transcriptomic profile may reflect elements of accelerated aging in OPFR-exposed animals, which is often related to cognitive impairment and anxiety disorders [101]. Unlike other estrogen-like EDCs such as Bisphenol-A, perinatal OPFR treatment did not affect estrogen or oxytocin receptor expression in the hippocampus [102]. However, there was very little overlap in the genes differentially expressed between hormone states, regardless of sex.

### 4.5 OPFR Treatment Affects Gene Expression in Hormone State Dependent Profiles

The most striking pattern between hormone status groups is the similarity of both up- and downregulated top gene ontology terms caused by OPFR exposure between males and high steroid state females. While many of these GO terms are still among the top differentially expressed modules in low steroid state females, they are differentially expressed in opposite directions in low females compared to males and high-steroid females. Some of this pattern is reflected in behavioral similarity between males and high females, in that OPFR-exposed males displayed some cognitive impairments in Barnes maze tasks and OPFR-exposed high steroid state females displayed cognitive impairments in the SOR and Y-maze tasks, while low steroid state females did not show any behavioral differences. This is notable, since previous work has shown more behavioral dissimilarity between males and (pro)estrus females than with diestrus females in fear conditioning [103], though this is not always the case in other behaviors, such as OFT or NOR [104, 105]. Nevertheless, developmental OPFR exposure clearly influences sex differences both behaviorally and transcriptionally.

Patterns in specific gene occurrences in the top GO terms are mirrored between low and high ovarian hormone status females caused by OPFR exposure. A number of mitochondrial/NADH genes are in the most downregulated GO terms in low-steroid females yet are among the most upregulated GO terms in high-steroid females. This suggests a significant difference in cellular metabolism between estrus cycle stages in the hippocampus caused by perinatal OPFR exposure that may contribute to the behavioral differences observed in the high-steroid females. Mitochondria are responsive to estrogens through estrogen receptor α and β activity and direct mitochondrial genomic action [106]. OPFRs such as TDCPP have also been shown to impair progesterone production and mitochondrial activity [107]. However, this brain-endocrine-mitochondrial interaction of OPFRs remains understudied [108], and this complex interaction affecting gene expression and possible hippocampal activity requires further examination.

The only transcript with increased expression caused by OPFR exposure in both low and high ovarian hormone status states but not in males was *Tmem229b*. In human ovarian-derived tissues, *tmem229b* is significantly upregulated by estrogen treatment and downregulated by estrogen inhibition via Fulvestrant exposure [109]. *Tmem229b* has been implicated with Parkinson’s disease development [110] and is upregulated in cultures of hypothalamic neural-progenitor/stem cells exposed to excess glucocorticoids [111], demonstrating how this gene could be related to cognitive and neurodevelopmental disorders. However, there is limited direct evidence in its relationship with endocrine disrupting chemical effects, warranting further study.

Otherwise, most of the OPFR-exposure’s effect on expression of individual genes is highly distinct between estrogen level profiles. Interestingly, *crhr1*, corticotropin releasing hormone receptor 1 transcript, was decreased only in OPFR-exposed low steroid state females, despite this group showing no difference in anxiety- or cognitive-related behaviors. Typically, estradiol increases expression of *crh* [112] and concurrent increased *crhr1* expression and high estradiol presence are necessary for acquisition of stress memories in males and females [113], potentially reflecting the patterns of altered expression in our males and high steroid state females in other metrics. Another gene with notably distinct expression between hormone states was *atn1*, atrophin 1, which was higher in high females, but lower in low females. Maternal exposure to the anesthetic sevoflurane causes elevated fetal hippocampal *atn1* exposure which alters neurodevelopment [114]. *Atn1* has also been implicated in neurodegenerative diseases such as Huntington’s [115]. The fact that OPRF exposure in development causes significant swings in expression with estrous cycle in the present data warrants further study.

## 5. Conclusion

In conclusion, perinatal exposure to OPFRs has sex and ovarian hormone dependent consequences. The similarity in spatial-cognitive impairments observed in OPFR-treated males and high-steroid state females could reflect the similarity in the ensembles of OPFR-altered functional gene ontology modules in the hippocampus. However, low ovarian steroid state may be somewhat protective of these effects, since low-steroid females had little observable cognitive impairment and opposing gene ontology ensembles. This may also be related to the significant OPFR-induced shift in gene ontology modules involving mitochondrial processes between low-and high-steroid females, with OPFR-induced downregulation of cellular metabolic processes in low-steroid females and OPFR-induced upregulation in high-steroid females. Similarly, there were opposing profiles of anxiety-like behavior between low- and high-steroid females exposed to OPFRs. This also aligns with decreased hippocampal transcript of crhr1 in OPFR-exposed low-steroid females, which “rebounds” to control levels in high-steroid state subjects. Perinatal OPFR exposure also has sex-associated effects, impairing males’ long-term memory, reducing hippocampal dopamine concentration, and upregulating expression of a functional hippocampal gene module related to extracellular matrix organization. These data support previous findings showing developmental OPFR exposure diminishes hippocampal formation in males, and suggests additional mechanisms involved in this reorganization. While the mechanisms that relate these shifts in cellular processes require further study, it is clear that perinatal exposure to OPFRs alters the hippocampus and hippocampus-related behaviors in a sex- and hormone-state dependent manner.

## Acknowledgments

The authors wish to thank the numerous undergraduate students who assisted with the perinatal dosing of the organophosphate flame retardants.

## Funding Sources

Supported by funds from the National Institutes of Health (R01MH123544; R21ES035889; P30ES005022). K.W. was previously supported by the National Institute of Environmental Health Sciences (T32ES007148; K99ES033256). K.W. is currently supported by the National Institute of Environmental Health Sciences (4R00ES033256). K.M. is currently supported by the National Institute of Environmental Health Sciences (T32ES007148).

## Author Contribution (CRediT)

K.W. Conceptualization, Data curation, Formal analysis, Funding acquisition, Investigation, Methodology, Project administration, Resources, Supervision, Validation, Visualization, Writing – original draft, Writing – review and editing

K.M. Data curation, Formal analysis, Methodology, Validation, Visualization, Writing – original draft, Writing – review and editing

R.M. Data curation, Formal analysis, Investigation, Writing – review and editing

J.E. Investigation, Writing – review and editing

V.A Investigation, Writing – review and editing

C.R. Investigation, Resources

A.Y. Investigation, Writing – review and editing

T.D. Investigation, Writing – review and editing

N.K. Investigation

T.A.R. Conceptualization, Funding acquisition, Project administration, Resources, Supervision, Writing – review and editing

## Data Availability

The data that support the findings of this study are openly available in Series GSE312005 at https://www.ncbi.nlm.nih.gov/geo/query/acc.cgi?acc=GSE312005. Raw data will be released early on request of reviewers and after publication. Analysis pipeline github repository: https://github.com/kmoran10/Project-OPFR-HPC-RNAseq.

## Notes

https://github.com/kmoran10/Project-OPFR-HPC-RNAseq

https://www.ncbi.nlm.nih.gov/geo/query/acc.cgi?acc=GSE312005

### Competing Interest Statement

The authors have declared no competing interest.

