## Supplementary figures and images for "Perinatal Exposure to Organophosphate Flame Retardants Induces Sex- and Hormone-Dependent Alterations in Anxiety, Memory, Neurotransmitter Content, and Hippocampal Gene Expression"

### supplemental figure 1

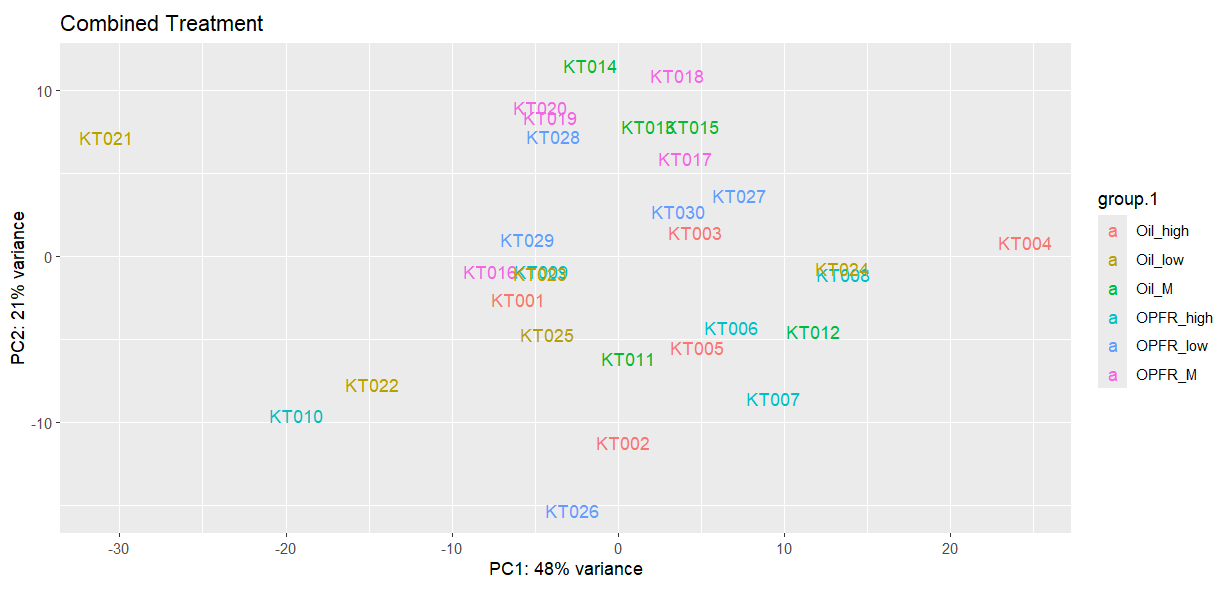

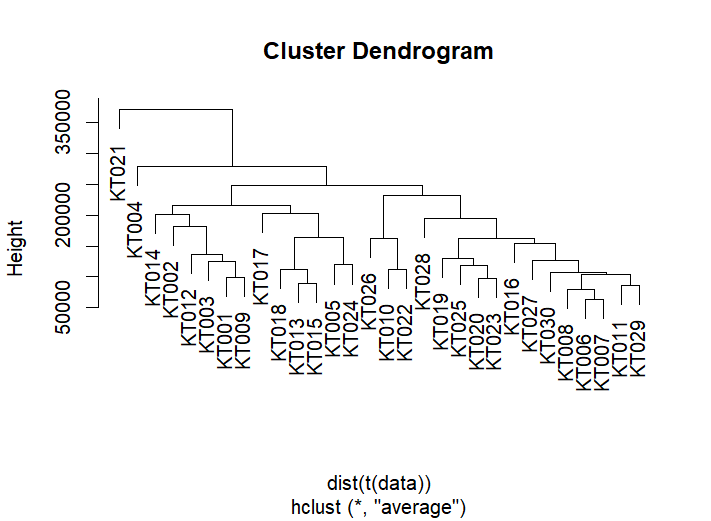
