## supplemental table 1 for "Perinatal Exposure to Organophosphate Flame Retardants Induces Sex- and Hormone-Dependent Alterations in Anxiety, Memory, Neurotransmitter Content, and Hippocampal Gene Expression"

| **Supplemental Table 1. Top DEGs across all hormone status groups** | | | | | | | |
| --- | --- | --- | --- | --- | --- | --- | --- |
| **ST1A. Low Ovarian Hormone Status Females** | | | | | | | |
| **gene** | **logFC** | **P.Value** | | | **chr** | | **description** |
| Zbtb39 | 0.246376 | <0.0001 | | | 10 | | zinc finger and BTB domain containing 39 |
| Crhr1 | -0.50362 | <0.0001 | | | 11 | | corticotropin releasing hormone receptor 1 |
| Pola2 | 0.263982 | <0.0001 | | | 19 | | polymerase (DNA directed), alpha 2 |
| Tle4 | 0.235037 | <0.0001 | | | 19 | | transducin-like enhancer of split 4 |
| Zfp719 | 0.288016 | <0.0001 | | | 7 | | zinc finger protein 719 |
| Epha7 | 0.219242 | <0.0001 | | | 4 | | Eph receptor A7 |
| Nol4 | 0.220278 | <0.0001 | | | 18 | | nucleolar protein 4 |
| Card10 | -0.28887 | <0.0001 | | | 15 | | caspase recruitment domain family, member 10 |
| Rtl4 | 0.345686 | <0.0001 | | | X | | retrotransposon Gag like 4 |
| Marcks | 0.240132 | <0.0001 | | | 10 | | myristoylated alanine rich protein kinase C substrate |
| Tob1 | 0.262038 | <0.0001 | | | 11 | | transducer of ErbB-2.1 |
| Phlda1 | 0.292386 | <0.0001 | | | 10 | | pleckstrin homology like domain, family A, member 1 |
| Golph3 | 0.211561 | <0.0001 | | | 15 | | golgi phosphoprotein 3 |
| Amer3 | 0.222943 | <0.0001 | | | 1 | | APC membrane recruitment 3 |
| Atp7a | 0.282863 | <0.0001 | | | X | | ATPase, Cu++ transporting, alpha polypeptide |
| Klhl9 | 0.20231 | <0.0001 | | | 4 | | kelch-like 9 |
| Rrp9 | -0.26702 | <0.0001 | | | 9 | | ribosomal RNA processing 9, U3 small nucleolar RNA binding protein |
| Mex3b | 0.344591 | <0.0001 | | | 7 | | mex3 RNA binding family member B |
| Gpr137b | -0.35452 | <0.0001 | | | 13 | | G protein-coupled receptor 137B |
| Zfp804a | 0.356622 | <0.0001 | | | 2 | | zinc finger protein 804A |
| Calhm5 | 0.466023 | <0.0001 | | | 10 | | calcium homeostasis modulator family member 5 |
| Trib2 | 0.287928 | <0.0001 | | | 12 | | tribbles pseudokinase 2 |
| Tmem229b | 0.268682 | <0.0001 | | | 12 | | transmembrane protein 229B |
| Snhg16 | -0.83692 | 0.0056 | | | 11 | | small nucleolar RNA host gene 16 |
| Pdik1l | 0.229772 | 0.0056 | | | 4 | | PDLIM1 interacting kinase 1 like |
| **ST1B. High Ovarian Hormone Status Females** | | | | | | | |
| **gene** | **logFC** | **P.Value** | | **chr** | | **description** | |
| Atn1 | -0.23708 | <0.0001 | | 6 | | atrophin 1 | |
| Ankrd35 | 0.51385 | <0.0001 | | 3 | | ankyrin repeat domain 35 | |
| Rita1 | 0.305 | <0.0001 | | 5 | | RBPJ interacting and tubulin associated 1 | |
| Gucd1 | 0.261204 | <0.0001 | | 10 | | guanylyl cyclase domain containing 1 | |
| Lias | 0.205811 | <0.0001 | | 5 | | lipoic acid synthetase | |
| Rasgrf1 | -0.25314 | <0.0001 | | 9 | | RAS protein-specific guanine nucleotide-releasing factor 1 | |
| Acvr1 | -0.31973 | <0.0001 | | 2 | | activin A receptor, type 1 | |
| Ankrd39 | 0.279178 | <0.0001 | | 1 | | ankyrin repeat domain 39 | |
| Mbnl1 | -0.23815 | <0.0001 | | 3 | | muscleblind like splicing factor 1 | |
| Qsox1 | 0.312646 | <0.0001 | | 1 | | quiescin Q6 sulfhydryl oxidase 1 | |
| Arl4d | 0.454877 | <0.0001 | | 11 | | ADP-ribosylation factor-like 4D | |
| Bop1 | 0.256049 | <0.0001 | | 15 | | block of proliferation 1 | |
| Dnajc21 | -0.39001 | <0.0001 | | 15 | | DnaJ heat shock protein family (Hsp40) member C21 | |
| Gorasp1 | 0.301548 | <0.0001 | | 9 | | golgi reassembly stacking protein 1 | |
| Gpbp1l1 | -0.20359 | <0.0001 | | 4 | | GC-rich promoter binding protein 1-like 1 | |
| Mogs | 0.227193 | <0.0001 | | 6 | | mannosyl-oligosaccharide glucosidase | |
| Nr2e1 | -0.24367 | <0.0001 | | 10 | | nuclear receptor subfamily 2, group E, member 1 | |
| Adipor2 | 0.263823 | <0.0001 | | 6 | | adiponectin receptor 2 | |
| Ascl1 | -0.43109 | 0.0028 | | 10 | | achaete-scute family bHLH transcription factor 1 | |
| Homer3 | 0.312117 | 0.0028 | | 8 | | homer scaffolding protein 3 | |
| Clasp1 | -0.2657 | 0.0028 | | 1 | | CLIP associating protein 1 | |
| L1cam | -0.27897 | 0.0028 | | X | | L1 cell adhesion molecule | |
| Kcnk4 | 0.233696 | 0.0028 | | 19 | | potassium channel, subfamily K, member 4 | |
| Sh2b2 | 0.624683 | 0.0028 | | 5 | | SH2B adaptor protein 2 | |
| Pepd | 0.275113 | 0.0028 | | 7 | | peptidase D | |
| **ST1C. Males** | | | | | | | |
| **gene** | **logFC** | **P.Value** | **chr** | | **description** | | |
| Irf7 | -0.87537 | <0.0001 | 7 | | interferon regulatory factor 7 | | |
| Dus4l | 0.363289 | <0.0001 | 12 | | dihydrouridine synthase 4-like (S. cerevisiae) | | |
| Slc13a5 | 0.242833 | <0.0001 | 11 | | solute carrier family 13 (sodium-dependent citrate transporter), member 5 | | |
| Rasip1 | -0.32638 | <0.0001 | 7 | | Ras interacting protein 1 | | |
| Robo4 | -0.46891 | 0.0052 | 9 | | roundabout guidance receptor 4 | | |
| Crem | 0.234615 | 0.0052 | 18 | | cAMP responsive element modulator | | |
| Abcg2 | -0.34734 | 0.0052 | 6 | | ATP binding cassette subfamily G member 2 (Junior blood group) | | |
| Tshz2 | -1.10126 | 0.0056 | 2 | | teashirt zinc finger family member 2 | | |
| Gask1a | 0.546412 | 0.0056 | 9 | | golgi associated kinase 1A | | |
| Gnb4 | -0.2072 | 0.0056 | 3 | | guanine nucleotide binding protein (G protein), beta 4 | | |
| Lin9 | 0.305347 | 0.0056 | 1 | | lin-9 DREAM MuvB core complex component | | |
| Poll | 0.296574 | 0.0056 | 19 | | polymerase (DNA directed), lambda | | |
| Myh11 | -0.47913 | 0.0064 | 16 | | myosin, heavy polypeptide 11, smooth muscle | | |
| Cdkl4 | -0.38627 | 0.0068 | 17 | | cyclin-dependent kinase-like 4 | | |
| Tti2 | 0.270935 | 0.0068 | 8 | | TELO2 interacting protein 2 | | |
| Fra10ac1 | 0.267472 | 0.0068 | 19 | | FRA10AC1 homolog (human) | | |
| Ccdc190 | 0.219609 | 0.008 | 1 | | coiled-coil domain containing 190 | | |
| Pinx1 | 0.401194 | 0.008 | 14 | | PIN2/TERF1 interacting, telomerase inhibitor 1 | | |
| Cldn10 | 0.22778 | 0.0088 | 14 | | claudin 10 | | |
| Sat1 | 0.340531 | 0.0088 | X | | spermidine/spermine N1-acetyl transferase 1 | | |
| Lnx1 | 0.20185 | 0.0104 | 5 | | ligand of numb-protein X 1 | | |
| Bex4 | 0.319672 | 0.0108 | X | | brain expressed X-linked 4 | | |
| Trim67 | -0.42229 | 0.0108 | 8 | | tripartite motif-containing 67 | | |
| Cdh5 | -0.45672 | 0.0108 | 8 | | cadherin 5 | | |
| Gm20754 | 0.337442 | 0.0108 | 3 | | predicted gene, 20754 | | |
